# An early Sox2-dependent gene expression program required for hippocampal dentate gyrus development

**DOI:** 10.1101/2020.08.16.252684

**Authors:** Sara Mercurio, Chiara Alberti, Linda Serra, Simone Meneghini, Jessica Bertolini, Pietro Berico, Andrea Becchetti, Silvia K. Nicolis

## Abstract

The hippocampus is a brain area central for cognition. Mutations in the human SOX2 transcription factor cause neurodevelopmental defects, leading to intellectual disability and seizures, together with hippocampal dysplasia. We generated an allelic series of Sox2 conditional mutations in mouse, deleting Sox2 at different developmental stages. Late Sox2 deletion (from E11.5, via Nestin-Cre) affects only postnatal hippocampal development; earlier deletion (from E10.5, Emx1-Cre) significantly reduces the dentate gyrus, and the earliest deletion (from E9.5, FoxG1-Cre) causes drastic abnormalities, with almost complete absence of the dentate gyrus. We identify a set of functionally interconnected genes (Gli3, Wnt3a, Cxcr4, p73 and Tbr2), known to play essential roles in hippocampal embryogenesis, which are downregulated in early Sox2 mutants, and (Gli3 and Cxcr4) directly controlled by SOX2; their downregulation provides plausible molecular mechanisms contributing to the defect. Electrophysiological studies of the Emx1Cre mouse model reveal altered excitatory transmission in CA1 and CA3 regions.

## Introduction

The hippocampus is a brain region important for cognition, playing essential roles in learning and in spatial and episodic memory formation. Hippocampus defects (of genetic origin, or acquired) can lead to intellectual disability (ID), deficits of memory formation, and epilepsy (Kandel, Schwartz, & Jessell, 2000).

Within the hippocampus, the dentate gyrus (DG) represents the primary input site for excitatory neuronal projections; the major type of DG neurons (granule neurons) are generated by neural stem cells (NSC) that are defined early in development, and continue neurogenesis during embryogenesis and also in postnatal stages, in mice as well as in humans (Berg et al., 2019; Zhong et al., 2020).

Patients carrying heterozygous loss-of-function mutations in the gene encoding the SOX2 transcription factor show a characteristic spectrum of central nervous system (CNS) defects, including hippocampal defects (involving the dentate gyrus), ID, and epilepsy (Fantes et al., 2003; Kondoh H, 2016; Ragge et al., 2005; Sisodiya et al., 2006). Understanding the developmental events and the genetic program controlled by SOX2 during hippocampal embryogenesis therefore provides a key to understand how their perturbation can lead to hippocampal disease (in SOX2-mutant patients and, more in general, in hippocampal defects of genetic origin).

In mouse, Sox2-dependent hippocampal disease has been previously modelled by conditional mutagenesis (Favaro et al., 2009). Sox2 pan-neural deletion at mid-embryogenesis, via a Nestin-Cre transgene, led to a relatively normal hippocampal development up to birth; at early postnatal stages, however, the hippocampus failed to complete its development, and remained hypoplastic, due to a failure of postnatal DG NSC. The study of SOX2 binding to DNA in NSC proved instrumental in the identification of various Sox2 target genes, playing important roles in the development of different brain regions in vivo, such as the basal ganglia (A. Ferri et al., 2013), the cerebellum (Cerrato et al., 2018), and the visual thalamus (Mercurio et al., 2019).

While postnatal hippocampal development was perturbed following Nestin-Cre-mediated Sox2 deletion, embryonic hippocampal development was, quite surprisingly, very little, if at all, affected in these mutants (Favaro et al., 2009). In principle, this could be due to redundant functions played by other homologous genes of the SoxB family, such as Sox1 and Sox3, coexpressed with Sox2 in the developing neural tube, and reported to function in hippocampal neural stem/progenitor cells (Rogers et al., 2013); alternatively, we reasoned that Sox2 may play non-redundant, very early functions in hippocampal development, that might not be revealed by Nestin-Cre-mediated deletion.

Here, we generated an allelic series of Sox2 conditional mutations, using Cre transgenes deleting Sox2 at stages earlier than Nestin-Cre: FoxG1-Cre, active from embryonic day (E) 8.5 (Hébert & McConnell, 2000), and Emx1-Cre (Gorski et al., 2002), active from E10.5. We report that early Sox2 deletion leads to drastic defects of hippocampal development, the earlier the deletion, the stronger the phenotype: in Emx1-Cre mutants, hippocampal development is perturbed, but still present, but in FoxG1-Cre mutants, hippocampal development is severely impaired, and the DG essentially fails to develop. We propose that Sox2 sets in motion a very early gene expression program in the hippocampal primordium, required for all of its subsequent development. Indeed, we show that early (but not late) Sox2 deletion reduces the expression of several genes (some of which SOX2-bound), individually characterized by previous studies as master regulators of hippocampal development (and human neurodevelopmental disease), including Gli3, Wnt3a, Cxcr4, Tbr2 and p73, some of which are known to cross-regulate each other.

## Results

### Sox2 is expressed in the primordium of the developing hippocampus and in the adjacent cortical hem

The transcription factor Sox2 is expressed throughout the neural tube from the beginning of its development (Avilion et al., 2003; Favaro et al., 2009; A. L. Ferri et al., 2004; Mariani et al., 2012). The hippocampus starts to develop around embryonic day (E) 12.5, in the medial wall of the telencephalon, and becomes morphologically recognizable in the following days (Fig. 1A) (Berg et al., 2019; Hodge et al, 2012). A region essential for the formation of the hippocampus is the cortical hem (CH), also known as the hippocampal organizer, identified in mice at E12.5; signaling from the CH is able to organize the surrounding tissue into a hippocampus (Grove, 2008; Mangale et al., 2008). The dentate neural epithelium (DNE), adjacent to the cortical hem (Fig. 1A), contains neural stem cells (NSC), that will generate granule neurons in the hippocampus dentate gyrus (DG) throughout development and, subsequently, in postnatal life (Berg et al., 2019). On the outer side of the neuroepithelium, towards the pia, a population of neurons, called Cajal-Retzius cells (CRC) (Fig. 1A) develops, that will have a key role in the morphogenesis of the hippocampus. NSC and intermediate neural progenitors (INP) will migrate from the DNE, along the dorsal migratory stream (DMS), towards the forming hippocampal fissure (HF), a folding of the meninges that will be invaded by CRC (Fig. 1A).

**Figure 1.**
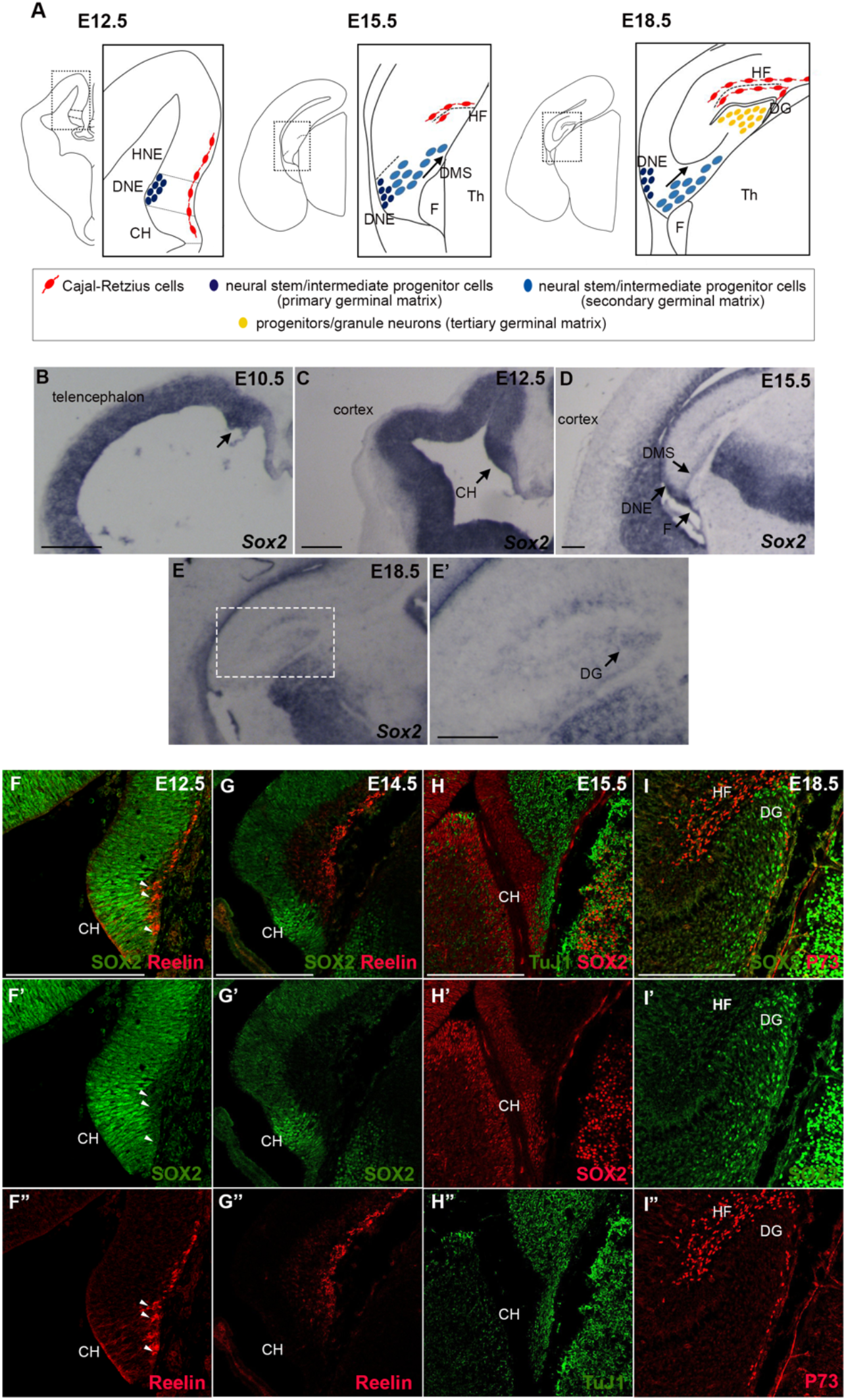
Sox2 expression in the dorsal telencephalon. **(A)** Schematic representation of the development of the hippocampus in the dorsal telencephalon. **(B-E)** *In situ* hybridization for *Sox2* on coronal section of mouse brains at E10.5 (B) E12.5 (C), E15.5 (D), E18.5 (E). Arrows indicate Sox2 expression in the developing hippocampus in particular in the dorsal telencephalon in (B), in the cortical hem (CH) in (C), in the dorsal migratory stream (DMS) in (D) and in the dentate gyrus (DG) in (E’). (**F-I)** Immunofluorescence of Sox2 (F-I), of markers of CRC, Reelin (F,G) and P73 (I), and of a marker of differentiating neurons TuJ1 (H). Representative single optical confocal sections are shown. Scale bars 200 μm. CH, cortical hem; DNE, dentate neuroepithelium; HNE, hippocampal neuroepithelium; DMS, dentate migratory stream; HF, hippocampal fissure; DG, dentate gyrus; F,fimbria; Th, thalamus.

We examined Sox2 expression by *in situ* hybridization (ISH) and immunofluorescence (IF), in the medial telencephalon, from which the hippocampus develops, between E12.5 and E18.5 (Fig. 1B-I). At E10.5 *Sox2* is expressed in the whole telencephalon including the dorso-medial region that will give rise to the hippocampus (Fig. 1B). At E12.5, Sox2 is expressed throughout the neuroepithelium in the medial telencephalic wall and it is enriched in the CH region (Fig. 1C); at E15.5, expression persists in the neuroepithelium, and is detected in the DMS and in the fimbria (a CH derivative) (Fig. 1D). Just before birth, at E18.5, Sox2 expression is detected in the developing DG (Fig. 1E,E’).

We then performed co-immunohistochemistry experiments with antibodies against SOX2, and markers of more differentiated cell types: CR cells markers Reelin and P73 (Fig. 1F,G,I) and the pan-neuronal marker TuJ1 (Fig. 1H). While SOX2 was detected in all cells within the neuroepithelium, as expected, we detected no or very little (Fig. 1F arrowheads) overlap with TuJ1, Reelin, or p73 (Fig. 1 F-I). Moreover, to test if Sox2 is expressed in the progenitors of CRC, we turned on EYFP in Sox2-expressing cells of the early telencephalon before CRC differentiation started, at E9.5 (via a Sox2-CreERT2 transgene and a lox-stop-lox reporter of Cre activity, Fig. S1), and found that these cells differentiated into Reelin-expressing CRC in the hippocampal fissure and the cortex (Fig. S1).

Thus, Sox2 expression in the developing hippocampus and CH is present mainly in undifferentiated neuroepithelial cells (including CRC precursors), and becomes extinguished in differentiation.

### Sox2 early ablation (FoxG1-Cre) prevents the development of the hippocampal dentate gyrus, and severely compromises hippocampal embryogenesis

Sox2 is required for postnatal development of the hippocampus, in particular to maintain NSC in the DG (Favaro et al., 2009); however, whether Sox2 has a role in hippocampus embryogenesis was not known. To address this question, we generated three different conditional knock-outs, to ablate Sox2 at different time points of telencephalon development. Specifically, we crossed a Sox2 floxed allele (Favaro et al., 2009) with the following Cre lines: FoxG1-Cre, deleting between E8.5 and E9.5 (A. Ferri et al., 2013; Hébert & McConnell, 2000), Emx1-Cre, deleting after E10.5 (Gorski et al., 2002), and Nestin-Cre, deleting after E11.5 (Tronche et al., 1999). The resulting conditional knock-outs (Sox2^flox/flox^;FoxG1-Cre; Sox2^flox/flox^;Emx1-Cre; Sox2^flox/flox^;Nestin-Cre) will be called FoxG1-Cre cKO, Emx1-Cre cKO and Nestin-Cre cKO respectively, from now onwards. As expected, complete Sox2 deletion is observed already at E9.5 in FoxG1-Cre cKO (in the whole telencephalon), and at E10.5 in Emx1-Cre cKO (in the dorsal telencephalon); in the Nestin-Cre cKO, deletion occurs after E11.5 (Favaro et al., 2009, Ferri et al., 2013, and data not shown).

We initially explored hippocampus development in the different mutants at the end of gestation (E18.5; P0), performing ISH with probes identifying hippocampal structures and cell types (Fig. 2). ISH for a general marker of the developing hippocampus, Cadherin 8 (Korematsu & Redies, 1997), shows that, at the end of gestation (E18.5, P0) the DG appears little, if at all, affected in the Nestin-Cre cKO (Fig. 2C), as expected (Favaro et al., 2009). However, in the Emx1-Cre cKO, the DG is greatly reduced, in particular anteriorly (Fig. 2B); remarkably, in the FoxG1-Cre cKO, the DG appears to be almost absent (Fig. 2A).

**Figure 2.**
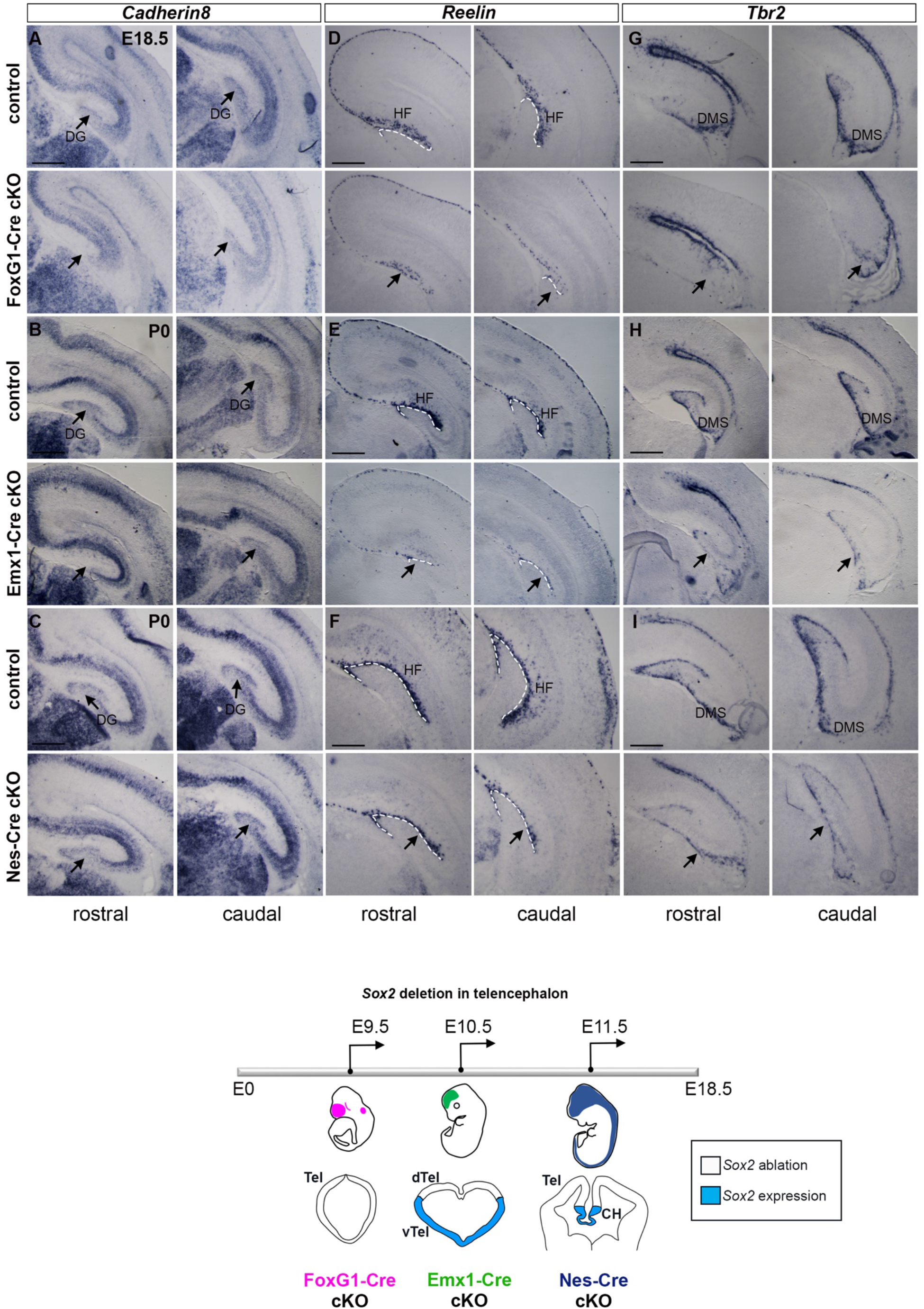
Hippocampus development is affected by Sox2 loss, the early Sox2 is ablated the stronger the phenotype observed. *In situ* hybridization for *Cadherin8* **(A-C)**, *Reelin* **(D-F)** and *Tbr2* **(G-I)** on coronal sections of control and Sox2 FoxG1-Cre cKO brains at E18.5 (A,D,G), control and Emx1-Cre cKO brains at P0 (B,E,H) and control and Nes-Cre cKO brains at P0 (C,F,I). At least 3 controls and 3 mutants were analyzed for each probe. A schematic representation of the timing of Sox2 ablation with the different Cre lines is at the bottom. Scale bars 200 μm. DG, dentate gyrus; HF, hippocampal fissure; DMS, dentate migratory stream; Tel, telencephalon; dTel, dorsal telencephalon; vTel, ventral telencephalon; CH, cortical hem.

At the end of gestation, CRC, expressing Reelin (D’Arcangelo et al., 1995), and INP, expressing Tbr2 (Hodge et al., 2013), have a characteristic organization in the hippocampus: CRC are localized around the HF, while INP have migrated from the DNE, by the ventricle, along the DMS, have reached the HF and are found below the CRC layer (see Fig. 1A). In the FoxG1-Cre cKO, Reelin expression (marking CRC) is greatly reduced, and a HF is not observed (Fig. 2D); in Emx1-Cre cKO, Reelin is reduced, but the HF is visible (Fig. 2E), and in Nestin-Cre cKO Reelin appears slightly reduced, but with a normal-looking distribution around the HF (Fig. 2F). Similarly, Tbr2 expression is greatly reduced in FoxG1-Cre cKO; an initial dorsal migratory stream is visible, but no DG is observed (Fig. 2G). Instead, in Emx1-Cre cKO, Tbr2-positive INP have reached the HF, but their abundance is greatly reduced (Fig. 2H). On the other hand, in Nestin-Cre cKO, Tbr2-positive INP appear to have completed their migration, and their abundance seems only slightly, if at all, reduced (Fig. 2I).

To summarize, Sox2 ablation by E9.5 in the telencephalon in FoxG1-Cre cKO results, by the end of gestation, in lack of DG formation, accompanied by a missing HF. Ablation just a day later, in Emx1-Cre cKO, has much less dramatic effects: a hippocampal fissure forms, though CRC and INP are reduced and the DG is much smaller compared to controls. Nestin-Cre cKO appear much less, if at all, affected, as previously published (Favaro et al., 2009).

### Neural progenitors, differentiated neurons and radial glia are affected by Sox2 loss

We then characterized the development of specific hippocampal cell types in the most affected (FoxG1-Cre) mutants. Key for the morphogenesis of the hippocampus is the radial glia (RG) scaffold known to be required for the DMS to reach its final destination in the forming DG (Li, Kataoka, Coughlin, & Pleasure, 2009). By immunohistochemistry for GFAP, recognizing RG, at E18.5, we find that the RG scaffold in FoxG1-Cre cKO is completely disorganized (Fig. 3A). No morphologically identifiable DG is present, and the few RG found have random organization (Fig. 3A, arrows). At this same stage, different neuronal populations are normally found in the hippocampus: granule neurons in the DG, and pyramidal neurons forming the CA1, CA2 and CA3 regions. We performed ISH for NeuroD1, a marker of differentiated neurons; in FoxG1-Cre cKO, while NeuroD1-positive cells in the CA regions are present, NeuroD1-positive cells in the DG, abundant in controls, are almost absent in the mutant (Fig. 3B).

**Figure 3.**
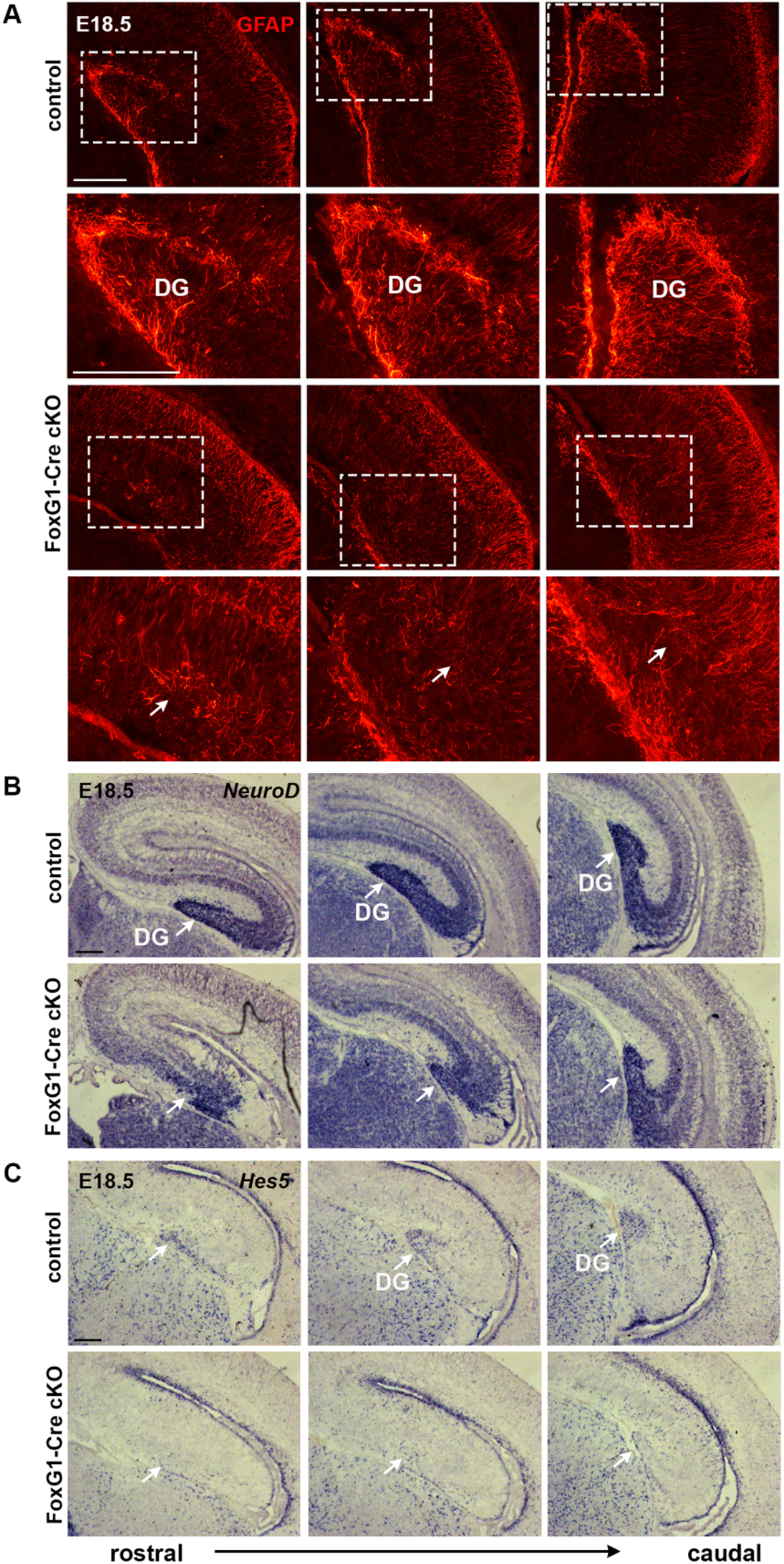
Radial glia formation, neuronal differentiation and neural stem cells formation is severely affected in FoxG1-Cre cKO. **(A)** GFAP immunofluorescence at E18.5 on coronal sections of control and FoxG1-Cre cKO hippocampi (controls n=6, mutants n=4). **(B,C)** *In situ* hybridization at E18.5 for *NeuroD* (B) (controls n=4, mutants n=3) and *Hes5* (C) (controls n=2, mutants n=2) on coronal sections of control and FoxG1-Cre cKO hippocampi. Arrows indicate the underdeveloped dentate gyrus (DG) in cKO. Scale bars 200 μm.

In the DG, at this stage, neural stem/progenitor cells, marked by the expression of the Hes5 gene (Basak & Taylor, 2007), are normally present (see Fig. 3C, controls); in FoxG1-Cre cKO, however, very few Hes5-positive cells are found (Fig. 3C).

In conclusion, early Sox2 loss in the telencephalon (FoxG1-Cre cKO) appears to lead to later reduction of both differentiated neurons and proliferating neural progenitors; in addition, the radial glia scaffold is completely disorganized.

### The formation of the hippocampal fissure and the dentate migration require Sox2 expression from early developmental stages

After having identified the hippocampal defects present, in our mutants, at the end of gestation, we examined earlier developmental stages, to define the developmental history of the defects. We focused in particular on the FoxG1-Cre mutant, showing the most pronounced abnormalities (see Fig. 2).

A defect in distribution of CRC (marked by Reelin) and INP (marked by Tbr2) is apparent, at the end of gestation, in Sox2 FoxG1-Cre and Emx1-Cre cKO (Fig. 2D,E,G,H). What happens in the first steps of the development of the hippocampus to CRC and INP in these mutants? We addressed this question by ISH with markers for these cell types at early developmental stages, in FoxG1-Cre (Early) cKO embryos (Fig. 4). We also examined the expression of Cxcr4, a chemokine receptor expressed in INP and neuroblasts in the DMS and in CRC, and its ligand Cxcl12, expressed by the meninges, and required for the migration of INP and CRC (Berger, Li, Han, Paredes, & Pleasure, 2007; Borrell & Marin, 2006; Hodge et al., 2013; Li et al., 2009); we also examined P73, a P53 homolog, expressed by CRC and important for hippocampal fissure and DG formation (Meyer et al., 2004; Meyer et al., 2019).

**Figure 4.**
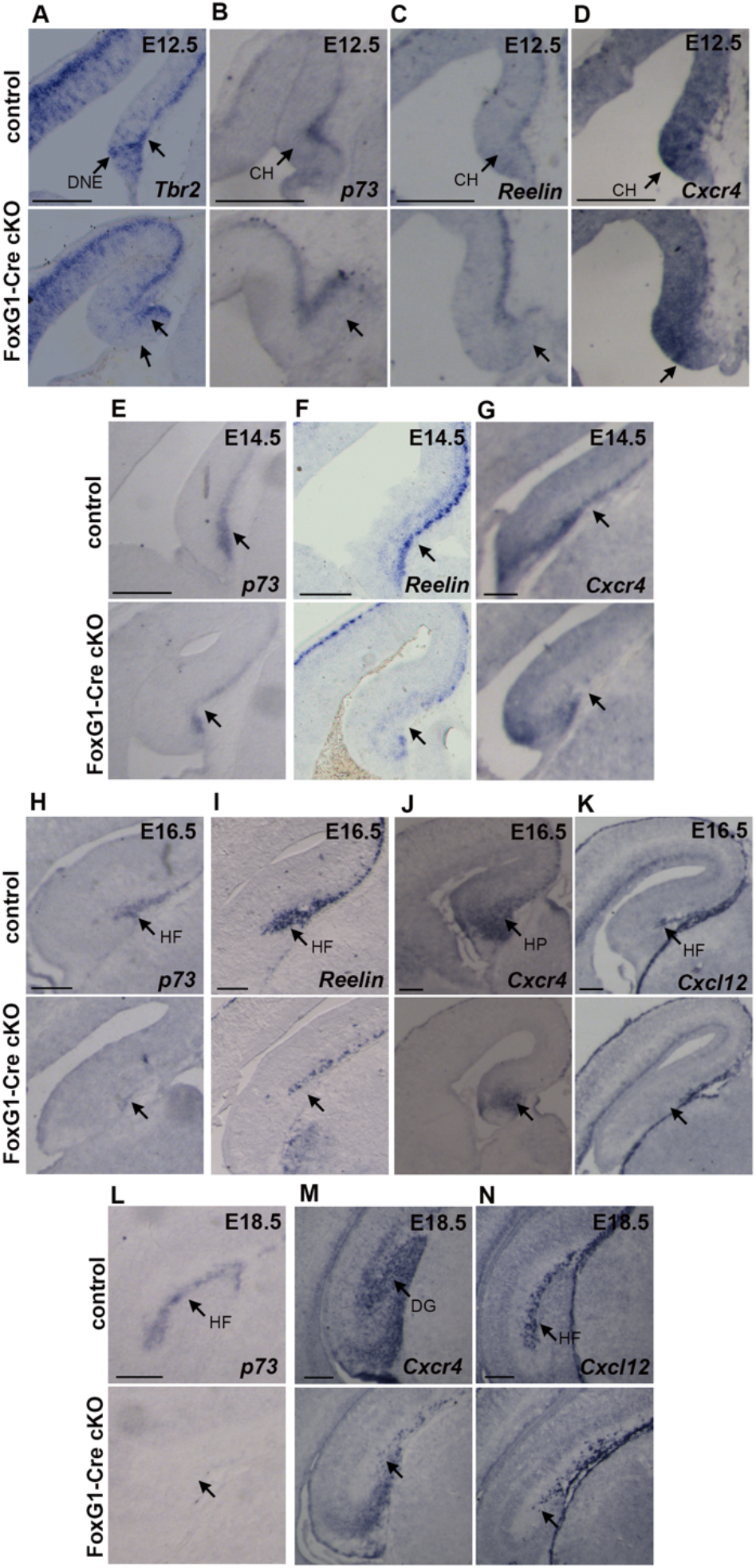
Expression of genes important for the development of the hippocampus is affected by Sox2 loss in FoxG1-Cre cKO. **(A-D)** *In situ* hybridization at E12.5 on coronal sections of control and FoxG1-Cre cKO dorsal telencephalons for *Tbr2* (controls n=10, mutants n=10) (A), *P73* (controls n=3, mutants n=3) (B), *Reelin* (controls n=7, mutants n=6) (C) and *Cxcr4* (controls n=7, mutants n=7) (D). **(E-G)** *In situ* hybridization at E14.5 on coronal sections of control and FoxG1-Cre cKO brains for *P73* (controls n=3, mutants n=3) (E), *Reelin* (controls n=7, mutants n=5) (F) and *Cxcr4* (controls n=3, mutants n=3) (G). **(H-K)** *In situ* hybridization at E16.5 on coronal sections of control and FoxG1-Cre cKO hippocampi for *P73* (controls n=2, mutants n=2) (H), *Reelin* (controls n=6, mutants n=5) (I), *Cxcr4* (controls n=5, mutants n=4) (J) and *Cxcl12* (controls n=4, mutants n=3) (K). **(L-N)**) *In situ* hybridization at E18.5 on coronal sections of control and FoxG1-Cre cKO hippocampi for *P73* (controls n=3, mutants n=3) (L), *Cxcr4* (controls n=5, mutants n=4) (M) and *Cxcl12* (controls n=5, mutants n=4) (N). Arrows indicate the downregulation of expression in the mutant cortical hem (CH), dentate neuroepithelium (DNE), hippocampal primordium (HP), dentate gyrus (DG) and hippocampal fissure (HF). Scale bars 200 μm.

At E12.5, Tbr2 is expressed, in controls, by INP in the DNE and in CR cells towards the pia (Fig. 4A); in the FoxG1-Cre cKO mutant, whereas Tbr2 expression in CRC (towards the pia, arrow) appears present, expression in the DNE is not detected (Fig. 4A). This might reflect a loss of Tbr2-expressing INP; however, we do not observe changes in the number of proliferating cells in this region at E12.5 by EdU labelling (Fig. S2), suggesting that at least some INP remain, but express less Tbr2 or are mislocalized. P73, Reelin and Cxcr4 expression appears unaltered in mutants compared to controls at this stage (Fig. 4B-D).

At E14.5, P73 and Reelin expression marks, in controls, CRC in the medial telencephalic wall region where hippocampal morphogenesis will soon begin (Fig. 4E,F, arrow); in the mutant, a strong reduction of P73 and Reelin expression is observed (Fig. 4E,F, arrow). Of note, this reduction is detected specifically in the CH of FoxG1-Cre cKO (Fig. 4E,F), even though Sox2 is ablated in the whole telencephalon. At this stage, also Cxcr4 expression in CRC appears reduced in the CH of FoxG1-Cre cKO (Fig. 4G).

At E16.5, in controls, strong P73 and Reelin expression marks the hippocampal fissure (HF), defining the beginning of overt hippocampal morphogenesis (Fig. 4H,I); in sharp contrast, this expression is not seen or greatly reduced in the mutant (Fig. 4H,I). Cxcl12 is also expressed, in the control, in the developing HF, and its expression is also lost in the mutant (Fig. 4K). Concomitantly, Cxcr4 expression in the hippocampus primordium (HP) is also reduced (Fig. 4J). These data point to a failure to initiate proper HF development in the mutant.

Interestingly, at 16.5, P73, Reelin and Cxcr4 expression is reduced throughout the telencephalon in FoxG1-Cre cKO (Fig. 4).

At E18.5, P73 marks the HF in controls, but its expression is completely absent in the FoxG1-Cre cKO brain, indicating a complete depletion of P73-positive CH-derived CRC (Fig. 4L). Cxcr4 expression in the DG and Cxcl12 expression in the HF is also greatly reduced in the mutant, confirming a severe abnormality of the mutant hippocampus at the end of gestation (Fig. 4M,N). In conclusion, the defects detected, at the end of gestation, in FoxG1-Cre mutants originate early in development, with a failure, at early stages, to develop a HF and migrating DNE cells in these mutants.

### Genes essential for hippocampal development are downregulated following early (FoxG1-Cre cKO), but not late (Emx1-Cre cKO, Nestin-Cre cKO), Sox2 deletion

Having observed that early Sox2 mutants (in particular FoxG1-Cre cKO) show severely defective hippocampal development, we searched for Sox2-regulated downstream genes, whose deregulation in mutants could explain the observed defects. We compared the expression of several candidate genes in mutants and controls, at E12,5, a stage preceding the observed abnormalities (clearly observed, in mutants, from E14.5, when hippocampal morphogenesis begins). Having observed that the defects in early Sox2 mutants (FoxG1-Cre cKO) are much more severe than those arising in later (Emx1-Cre and Nestin-Cre cKO) mutants, we reasoned that genes downstream to Sox2, that are functionally relevant for these early defects, should show altered expression in early (FoxG1-Cre) mutants, but not, or less, in later mutants (Emx1-Cre; Nestin-Cre).

We thus investigated the expression of genes, representing candidate mediators of Sox2 function, in early and late mutants, by ISH.

Prime candidate genes to mediate defective hippocampal development in early Sox2 mutants include genes encoding signalling molecules, expressed in the CH.

Key signaling molecules secreted by the CH and required for hippocampus formation are components of the Wnt pathway; in fact, Wnt3a knock-out results in a complete loss of the hippocampus (Lee, Tole, Grove, & McMahon, 2000). We analyzed what happens, at E12.5, to the expression of Wnt3A in the three Sox2 cKO. We found that Wnt3A is severely downregulated specifically in the CH of FoxG1-Cre cKO (Fig. 5A), but only slightly downregulated in Emx1-Cre cKO (Fig. 5B), while it is only very mildly, if at all, reduced at this stage in the Nestin-Cre cKO (Fig. 5C). We analyzed the expression of another Wnt family member, Wnt2b in FoxG1-Cre cKO and Emx1-Cre cKO. While Wnt2b was strongly downregulated in the CH of FoxG1-Cre cKO (Fig. 5D), it was only slightly downregulated in the CH of Emx1-Cre cKO compared to controls (Fig. 5E). Wnt5A, another Wnt family member normally expressed in the CH, was instead expressed in the CH of FoxG1-Cre cKO (Fig. 5F), indicating that the CH, as a structure, is present in these mutants, though it fails to express Wnt3a and Wnt2b. Interestingly, expression of the transcription factor Lhx2, a marker of the cortex which is not expressed in the cortical hem, has a normal expression pattern in FoxG1-Cre cKO, including an Lhx2-non-expressing neuroepithelial region, suggesting that a CH is present in these mutants (Fig. 5G).

**Figure 5.**
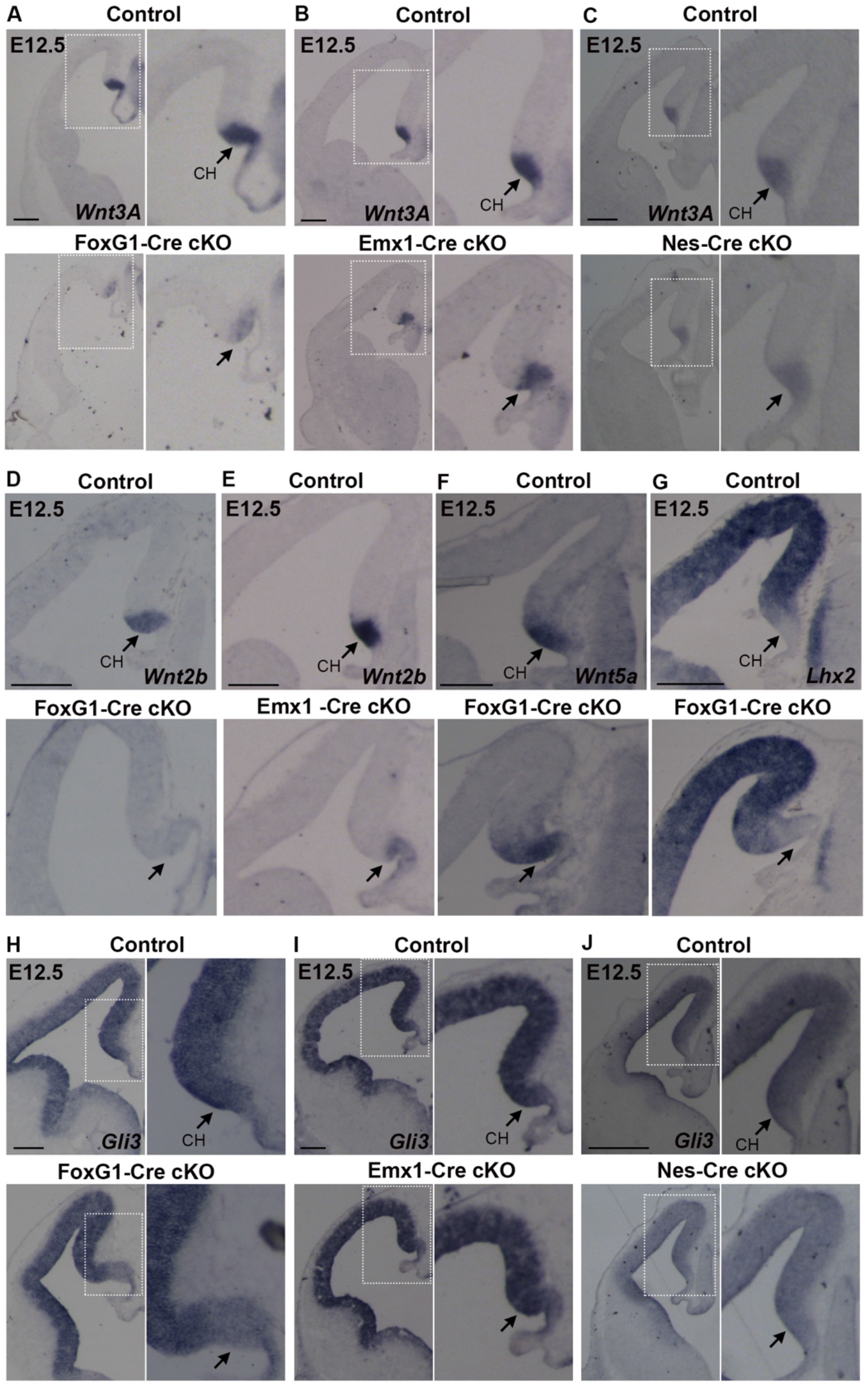
Expression of key molecules for hippocampal development is downregulated in the cortical hem of FoxG1-Cre cKO but mildly or not affected in Emx1-Cre or Nes-Cre cKO. *In situ* hybridization at E12.5 for Wnt3A **(A-C)**, *Wnt2b* **(D,E)**, *Wnt5A* **(F)**, *Lhx2* **(G)**, Gli3 **(H-J)** on control and FoxG1-Cre cKO (A,D,F,G,H), Emx1-Cre cKO (B,E,I) and Nes-Cre cKO (C,J) coronal brain sections. Arrows indicate the cortical hem (CH). At least 3 controls and 3 mutants were analyzed for each probe. Scale bars 200 μm.

In conclusion, expression of components of the Wnt pathway known to be involved in the development of the hippocampus is strongly downregulated in the CH of FoxG1-Cre cKO, but not of Emx1-Cre cKO and Nestin-Cre cKO.

Other key genes for hippocampus formation include Gli3, encoding a transcription factor acting as a nuclear effector in the Shh signaling pathway. The knockout of Gli3 impairs the development of the hippocampus, where DG development is as severely affected as in our Sox2 early (FoxG1-Cre cKO) mutants. Of note, Gli3 acts, in hippocampal development, by regulating expression of components of the Wnt pathway (Grove, Tole, Limon, Yip, & Ragsdale, 1998). We found that Gli3 expression is specifically downregulated in the CH (though not in the cortex) of FoxG1-Cre cKO, but not of Emx1-Cre cKO and Nestin-Cre cKO (Fig. 5H-J).

Recent work from our laboratory identified SOX2 binding sites in an intron of the Gli3 gene in NSC cultured from the mouse forebrain; further, this intronic region is connected to the Gli3 promoter by a long-range interaction mediated by RNApolII ((Bertolini et al., 2019) and Fig. 6A). A DNA segment, overlapping the SOX2 peak, drives expression of a lacZ transgene to the embryonic mouse forebrain ((Visel et al., 2009) and https://enhancer.lbl.gov) (Fig. 6A). We found that this Sox2-bound region, when connected to a minimal promoter and a luciferase reporter gene, and transfected in Neuro2a cells, is activated by increasing doses of a cotransfected Sox2-expressing vector in a dose-dependent way (Fig. 6B).

**Figure 6.**
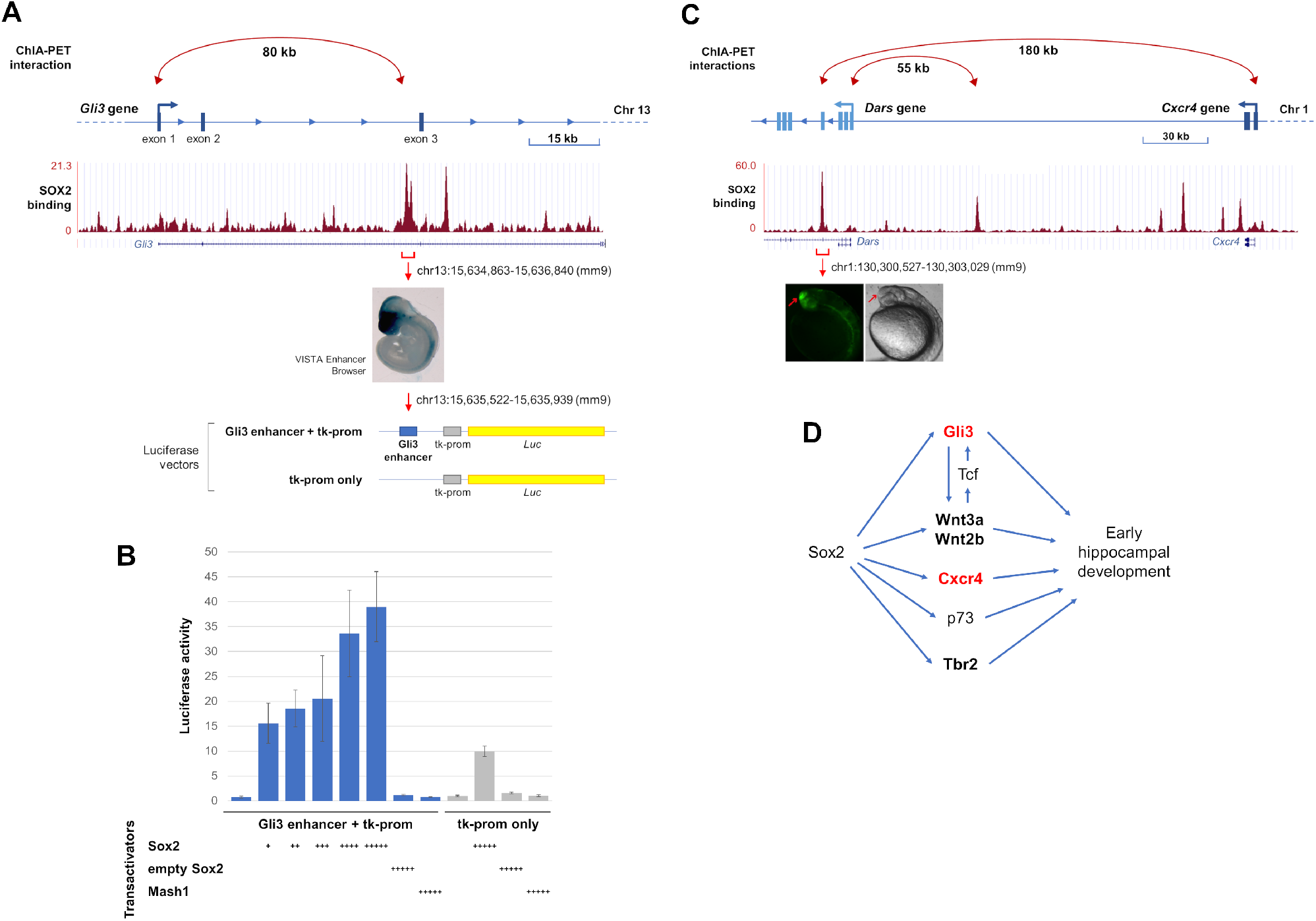
SOX2 acts on distal enhancers and on long-range enhancer-promoter interactions of several genes key to hippocampal development, and activates a Gli3 intronic enhancer in a dose-dependent way. (A) Diagram of the Gli3 gene, and SOX2-binding profile across the Gli3 locus in NSC (ChIPseq data from (Bertolini et al., 2019). A Sox2-dependent 80kb long-range interaction connects the Gli3 promoter with a SOX2-bound region, in the second intron (ChIA-PET data from (Bertolini et al., 2019). This region acts as a brain-specific enhancer in E10.5 mouse embryo (image from https://enhancer.lbl.gov/); it was cloned into the depicted luciferase vector, upstream to a minimal tk promoter, to address its responsivity to Sox2. (B) Enhancer activation assay in Neuro2a cells transfected with the constructs in (A): Gli3 enhancer + tk-promoter (blue histograms), or tk-promoter only (grey histograms). Cotransfection of these constructs with increasing amounts of a Sox2-expressing vector (Sox2, X axis), but not of a control “empty” vector (empty Sox2), or a Mash1-expressing vector (Mash1), resulted in dose-dependent increase of luciferase activity (Y axis) driven by the Gli3 enhancer + tk-prom vector, but not the tk-prom only vector. The molar ratios, compared with the luciferase vector (set at 1) were: +, 1:0.050; ++, 1:0.075; +++, 1:0.125; ++++, 1:0.25; +++++, 1:0.5. Results are represented as fold-change increase in activity compared with the tk-prom only vector, which is set at 1. Values are the mean of two (for Sox2+ and Sox2++) or three (other samples) independent experiments carried out in triplicate. Error bars represent standard deviation. (C) Diagram of the Cxcr4 gene, reporting SOX2 binding and Sox2-dependent long-range interactions in NSC (as in A for Gli3; data from (Bertolini et al., 2019)). Note that the Cxcr4 promoter is connected to a SOX2-bound region within the intron of a different gene, Dars; this region acts as a brain-specific enhancer in transgenic zebrafish embryos (picture from (Bertolini et al., 2019)). (D) A model depicting the activation, by Sox2, of different genes key to hippocampal development (present paper), some of which cross-regulate each other; in red, direct SOX2 targets; in bold, early expressed hippocampal regulators, downregulated already at early stages in Sox2 mutants (see Discussion).

Cxcr4, downregulated in early (FoxG1Cre) Sox2 mutants at E14.5 (Fig. 4G), is also functionally involved in the development of the hippocampus (Discussion). Of note, an enhancer active in the developing brain, located within an intron of the Dars gene, but connected to the Cxcr4 gene promoter by a long-range interaction in brain-derived NSC chromatin, is bound by SOX2 in these cells (Bertolini et al., 2019) (Fig. 6C).

In conclusion, Sox2 early ablation leads to reduced expression, particularly in early (FoxG1-Cre) mutants, of several genes key to hippocampal development, some of which are directly bound and regulated by SOX2; some of these genes (Gli3, Wnt3a) are also known to functionally regulate each other (see Discussion). These genes may thus be considered as part of a Sox2-dependent gene regulatory network, controlling hippocampal development (Fig. 6D; see Discussion).

### Emx1-Cre mediated Sox2 ablation alters the excitatory input in CA3 and CA1 pyramidal neurons

We also wished to ask about the consequences of Sox2 early loss on the physiological functioning of the postnatal hippocampus.

As illustrated earlier (Fig. 2A, 2B), early Sox2 loss causes DG hypoplasia, most severe in FoxG1-cKO mutants, but clearly present also in Emx1-Cre cKO mice. Since FoxG1-cKO are perinatally lethal (A. Ferri et al., 2013), we performed physiology studies on Emx1-cKO mutants. We addressed, in particular, the function of CA3 and CA1 pyramidal neurons, central to hippocampal circuitry and relatively spared, morphologically at least, in our mutants (in comparison to the severely hypoplastic DG).

The DG receives its main extrinsic input from the entorhinal cortex and is the first hippocampal station of the classical trisynaptic pathway: entorhinal cortex → DG granule cells → CA3 pyramidal neurons → CA1 pyramidal neurons. The DG projects exclusively to CA3 through mossy fibers. In turn, CA3 projects to CA1 through Schaffer collaterals (Witter & Amaral, 2004). Hence, we investigated whether the hypoplastic DG in our Emx1-Cre mutants could alter signal transfer to CA3 and CA1. This hypothesis was tested by studying intrinsic excitability and excitatory transmission in CA3/CA1 pyramidal neurons. These were first identified by their typically large pyramidally-shaped soma (∼20 μm diameter, in CA3), and then further distinguished by their action potential firing. We focused on regular-spiking pyramidal neurons, the widest population, characterized by slow firing with modest adaptation and excitability properties consistent with literature on CA1-CA3 neurons in mice (e.g., (Hunt, Linaro, Si, Romani, & Spruston, 2018; Venkatesan, Liu, & Goldfarb, 2014)). A typical example is shown in Fig. 7A. The excitability features of pyramidal neurons from control and mutant mice are shown in Supplementary Table 1, while the stimulus/frequency relations are shown in Fig. 7B. Overall, little difference was observed in intrinsic excitability between mutant and control mice, in both CA1 and CA3.

**Figure 7.**
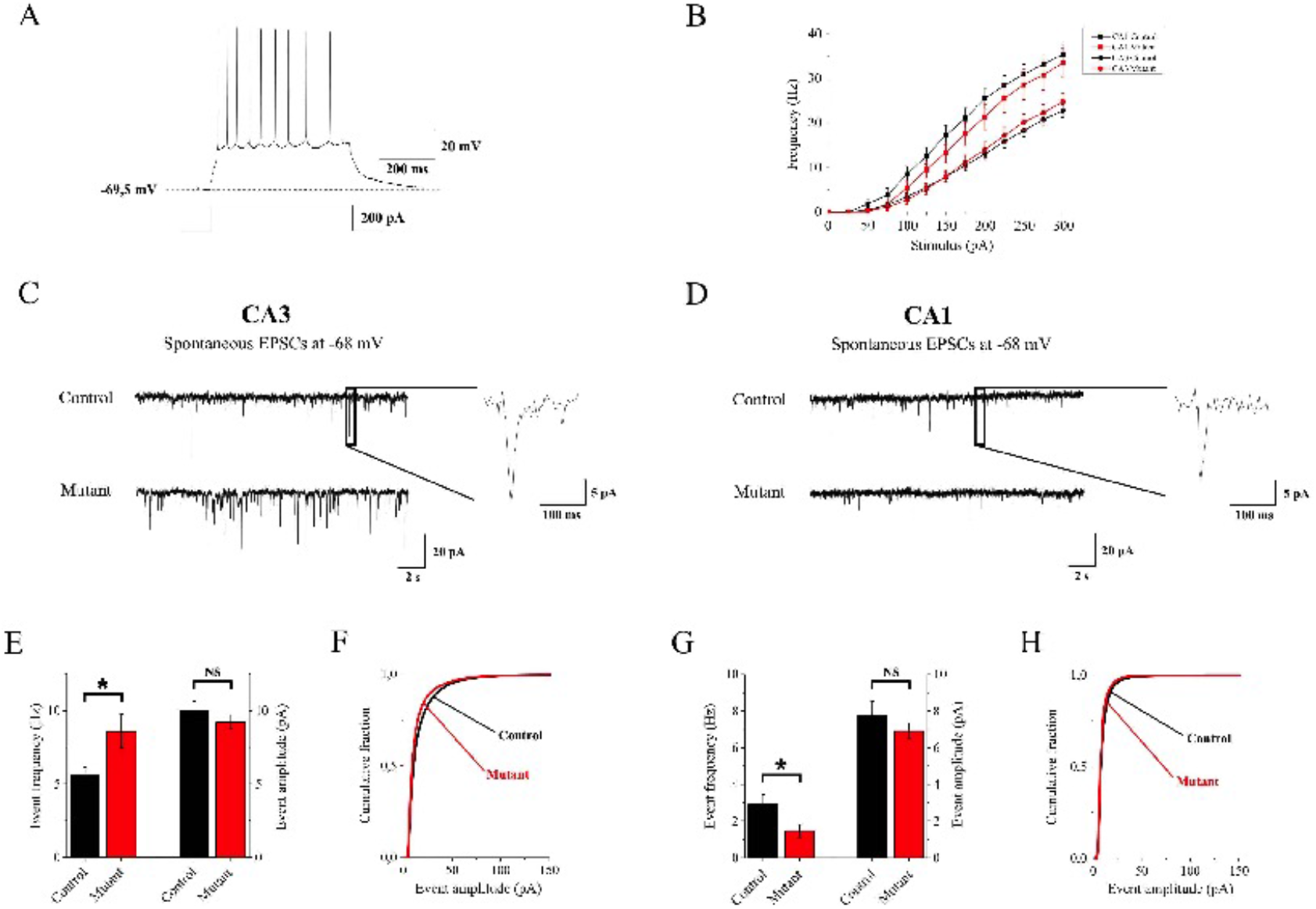
In Emx1-Cre cKO mice, excitatory transmission is altered in CA3 and CA1 hippocampal regions. Early Sox2 ablation leads to alterations in the excitatory input onto both CA3 and CA1 pyramidal neurons. (A) Typical firing response to a 200 pA stimulus of injected current in a CA3 pyramidal neuron. (B) Average stimulus/frequency relation for hippocampal pyramidal neurons recorded in CA3 (*Circles*) and CA1 (*Squares*). No major differences were observed between control (*Black*) and mutant animals (*Red*). (C and D) EPSCs traces, at −68 mV, recorded in simulated physiologic conditions onto pyramidal neurons in CA3 and CA1 region respectively. *Insets*. Magnification of a representative EPSC event. (E) Average EPSCs frequencies and median amplitudes observed in CA3 pyramidal neurons recorded from 15 animals between p19 and p31. In mutant animals, Sox2 ablation induced a significant increase in EPSCs frequency compared to controls (8.60 ± 1.15 Hz, n = 20 and 5.59 ± 0.57 Hz, n = 28 respectively; p = 0.03941, with Mann-Whitney test), whereas, no significant effect was produced on event amplitudes (9.21 ± 0.47 pA, n = 20 and 10.01 ± 0.61 pA, n = 28 respectively). (F) Amplitude distribution of the total amount of collected EPSCs showing no major differences between control and mutant mice. (G) In CA1, EPSCs frequency significantly decreased in mutant animals compared to controls (1.46 ± 0.35 Hz, n = 13 and 2.91 ± 0.52 Hz, n = 13 respectively; p = 0.02745, with Mann-Whitney test). No difference in the average median amplitude was observed (controls: 7.75 ± 0,69 pA, n = 20 and mutants: 6.92 ± 0.44 pA, n = 28). (H) The amplitude distribution of the total pool of events recorded from 13 animals between p19 and p31 showed no major alterations between control and mutant mice.

In these neurons, we recorded the spontaneous excitatory post-synaptic currents (EPSCs) for 10 min after reaching the whole-cell configuration, at −68 mV. Spontaneous EPSCs reflect the overall excitatory input impinging on a given pyramidal neuron. Typical EPSC traces from CA3 pyramidal neurons are shown in Fig. 7C, for controls and mutants. Somewhat surprisingly, EPSC frequencies in CA3 displayed a ∼30% increase in mutant animals compared to the controls (Fig. 7E). On the contrary, the average EPSC median amplitudes were not different between control and mutant mice (Fig. 7E). Moreover, the EPSC amplitudes obtained from all control and mutant cells were pooled in Fig. 7F. The amplitude distributions of the two genotypes were compared with KS test, which revealed no significant difference.

Next, we studied the excitatory input onto CA1, which is the last station of the hippocampal serial pathway of information transfer. Typical EPSC traces are shown in Fig. 7D for control and mutant. As expected (Traub, Jefferys, & Whittington, 1999), the overall EPSC frequency and amplitude tended to be smaller in CA1, compared to CA3. The average EPSC frequencies in CA1 are reported for control and mutant mice in Fig. 7G. Data reveal an approximately 50% reduction in mutant animals compared to the controls. Once again, little difference between genotypes was observed in the EPSC amplitudes (Fig. 7G, H).

In conclusion, CA3/CA1 pyramidal neuron firing or EPSC amplitudes were not altered in Emx1-Cre cKO mice, arguing against a direct effect of the mutation on the synaptic machinery or intrinsic excitability, which is consistent with the lack of expression of Sox2 in these neurons (data not shown). However, EPSC frequency increased in CA3 and was approximately halved in CA1 of mutant mice, suggesting that excitatory signal transfer along the canonical trisynaptic pathway was unbalanced as a consequence of the major impairment of DG development produced by early Sox2 ablation.

Overall, our data indicate significant functional alterations of the hippocampal circuitry in early Sox2 mutants, which might plausibly contribute to the epileptic and cognitive defects in human patients (see Discussion).

## Discussion

In this work, we highlighted an early time window in hippocampal development, where the Sox2 transcription factor is necessary to initiate the embryogenesis of the hippocampus. In fact, following Sox2 mutation with FoxG1-Cre, active from E8.5 (A. Ferri et al., 2013; Hébert & McConnell, 2000), hippocampal development is drastically defective, with a nearly complete absence of the DG; DG development is also defective, but present, following mutation with Emx1-Cre, active from E10.5; Sox2 mutation with Nestin-Cre has very little effect on hippocampal embryogenesis (Fig. 2).

These observations point to gene regulatory events, orchestrated by Sox2, that are required to initiate hippocampal development; at least some of these events are likely to be direct effects of SOX2 (Fig. 6). What is the nature of these events?

### Gene regulatory events mediating early Sox2 function in hippocampal development

A severe reduction in the early expression of key regulators of hippocampal development (Wnt3a; Gli3; Cxcr4; Tbr2; p73) is observed in early Sox2 mutants (FoxG1-Cre cKO), already at early stages of hippocampal embryogenesis (E12.5, E14.5), preceding the overt phenotypic manifestation of the defect (hippocampal morphogenesis begins at about E14.5) (Figs. 4,5). Of note, reduced expression in the mutant CH at E12.5 is seen for some genes (e.g. Wnt3a, Gli3) but not others (Wnt5a; Fig. 5), suggesting that the CH is present, but misfunctional in directing hippocampal formation. Importantly, a reduction in expression of these master genes is also observed in Emx1-Cre mutants, but to a lesser extent than in FoxG1-Cre mutants (Fig. 5). These observations suggest that the differential reduction in the expression of these key regulators accounts, at least in part, for the differences in the severity of the hippocampal embryogenesis defects between the three mutants.

What molecular mechanisms cause the differential expression of master hippocampal regulator genes between different mutants?

SOX2 is able to directly bind to at least some of these target genes (Gli3, Cxcr4, Fig. 6A,C) in neural cells chromatin, and to act as a transcriptional activator on some SOX2-bound enhancers (Gli3) within these loci (Fig. 6B), suggesting that it is directly involved in the transcriptional activation of at least some of these genes during hippocampal development. Further, SOX2-bound distant enhancers within the Gli3 and Cxcr4 loci are connected to the gene promoter in a Sox2-dependent way, at least in NSC (Fig. 6A,C), indicating that SOX2 may contribute to their regulation also through this “architectural” function (Bertolini et al., 2019; Wei, Nicolis, Zhu, & Pagin, 2019).

At E12.5 and afterwards, Sox2 is ablated in both FoxG1-Cre and Emx1-Cre mutants, yet critical genes are much more downregulated in FoxG1-Cre mutants, in agreement with a requirement for Sox2 to properly initiate the expression of these genes at early stages. We speculate that SOX2 may act at early stages to initiate the organization of a 3D interaction network connecting gene promoters to enhancers (Fig. 6A,C), as a prerequisite for gene expression, in agreement with previous findings in NSC (Bertolini et al., 2019; Wei et al., 2019).

### Altered regulation of a gene regulatory network of hippocampal master genes leads to defective cell development and cell-cell signaling in early Sox2 mutants, and eventually to defective hippocampal structure and function

The failure to properly activate early-acting hippocampal master genes may provide a molecular explanation to the failure to develop, in early Sox2 mutants, cell types essential in hippocampal development, or to prevent their proper behavior, as observed in Figs. 2,3.

In FoxG1-Cre, but not in Emx1-Cre Sox2 mutants, the Gli3 gene is downregulated at early stages (E12.5) in the segment of medial telencephalic wall required for hippocampal development, the CH (Fig. 5H,I). Gli3 encodes a transcription factor, and its homozygous mutation (as in the *extra-toes* Gli3^Xt/Xt^ mouse mutant) leads to absence of the hippocampus (Li & Pleasure, 2014; Theil, Alvarez-Bolado, Walter, & Ruther, 1999). In Gli3 mutant embryos, the medial wall of the telencephalon fails to invaginate to initiate hippocampal development, pointing to an early defect of the proliferating neuroepithelial cells of the prospective hippocampus (Li & Pleasure, 2014; Theil et al., 1999). Mutations of human GLI3 cause Pallister-Hall syndrome and Greig cephalopolysyndactyly syndrome, a complex defect that can involve seizures and intellectual disability, though hippocampal abnormalities were not specifically investigated (Naruse, Ueta, Sumino, Ogawa, & Ishikiriyama, 2010). In mouse Gli3 mutants, the expression of Wnt signaling molecules, normally expressed in the CH, including Wnt3a and Wnt2b, is lost, and Wnt signaling is impaired at early stages of hippocampal development (Fotaki, Price, & Mason, 2011; Grove et al., 1998; Theil, Aydin, Koch, Grotewold, & Ruther, 2002).

The expression of Wnt3a and Wnt2b, encoding secreted signaling molecules produced by the CH signaling center, is also defective in early Sox2 mutants (Fig. 5); their downregulation is most pronounced in FoxG1-Cre mutants, less so in Emx1-Cre mutants (see above and Fig. 5A,B,D,E). The downregulation of Gli3 may contribute to this (see above).

Wnt signaling exerts its effects on target cells by inducing nuclear translocation of beta-catenin, that acts as a transcriptional regulator associating with TCF transcription factors; mutation of TCF factors, e.g. Lef1, leads to failure of hippocampal development (Galceran, Miyashita-Lin, Devaney, Rubenstein, & Grosschedl, 2000; Li & Pleasure, 2014; Roelink, 2000). Of note, TCF binding regulates (Hasenpusch-Theil et al., 2012) the same intronic Gli3 enhancer, that we found to be bound and activated by Sox2 (Fig. 6A,B), suggesting that this element may integrate the effects of Wnt signaling and SOX2 activity in controlling Gli3 expression. Interestingly, Sox2/TCF binding sites were also described to act on other genes in the context of a transcriptional switch accompanying chromatin remodeling during neuronal differentiation (Muotri et al., 2010).

We attempted to reactivate the Wnt pathway in the FoxG1-Cre cKO, by LiCl injection, to see if we could rescue any of the observed defects. We found some amelioration of the organization and number of CRC in the cortex (Fig. S3B,C), although the overall hippocampus development remained defective (Fig. S3A,C). We also tried to reactivate the Wnt pathway by a Wnt agonist (AZD 1080); a partial rescue of Reelin retention in the CH, usually observed in mutants, was observed at E14.5 in the FoxG1-Cre cKO (Fig. S3D). We hypothesized that earlier treatment might have had more pronounced effects, however this resulted in high embryonic lethality, preventing us to observe the effects.

In conclusion, we propose that loss of Wnt signaling from the CH represents one mechanism whereby Sox2 early loss causes defective hippocampal embryogenesis likely by regulating the production of CRC. Assessing the relative contribution of this mechanism will be postponed to future studies.

We detected, in our early mutants, reduced expression of Tbr2, Cxcr4, Cxcl12 and p73, marking specific cell types in hippocampal embryogenesis (Fig. 4). However, knock-out experiments previously demonstrated that these genes, further to marking specific cell types (see Fig. 2), also play functional roles in hippocampal (as well as neocortical) development (Bagri et al., 2002; Hodge et al., 2013; Lu, Grove, & Miller, 2002; Meyer et al., 2004; Mimura-Yamamoto et al., 2017). This suggests that their reduced expression in Sox2 mutants may also functionally contribute to the hippocampal defects.

Cxcr4, whose expression is downregulated at early stages in Sox2 early mutants (Fig. 4G), is essential in particular for the development of the DG (Lu et al., 2002; Mimura-Yamamoto et al., 2017). Cxcr4 encodes a cell surface receptor, expressed in Granule Cell Progenitors (GCP) of the developing hippocampus, that also express GFAP (Mimura-Yamamoto et al., 2017). In hippocampal development, GCP, arising in the ventricular zone (DNE), migrate (dentate migratory stream) to the subpial region, to form the granule cell layer (GCL) of the DG (Fig. 1A). The production and migration of GCP is regulated by various signaling molecules, including CXCL12 (the CXCR4 ligand), Reelin, Wnt, and BMP proteins, secreted by regions surrounding the developing DG. In the absence of Cxcr4, the numbers of dividing cells in the migratory stream and the prospective DG is dramatically reduced (Lu et al., 2002). It thus seems plausible that Cxcr4 deficiency importantly contributes to the impaired development of GFAP-positive GCP, and the consequent failure to develop a DG, seen in our early Sox2 mutants.

P73 encodes a transcription factor expressed in differentiating CRC (Fig. 4), the choroid plexus and the ependyma (Meyer, Schaaps, Moreau, & Goffinet, 2000; Yang et al., 2000) and its knock-out in mice results in a phenotype very similar to the early loss of Sox2 in FoxG1-Cre cKO, with a lack of HF and almost absent DG (Meyer et al., 2019). P73 has a similar expression pattern in the fetal human brain suggesting a role in hippocampus development also in humans (Meyer et al., 2019). Interestingly, Reelin-expressing CRC in Sox2 mutants are similarly reduced in number and they may be retained in the cortical hem instead of moving towards the pia. P73 has a very restricted expression pattern, but its knock-out has a broad effect on cortical patterning suggesting it could be involved in the signaling activities of the CH (Meyer et al., 2004).

### Radial scaffold, Cajal-retzius cells and lack of hippocampal fissure and dentate gyrus

One of the key outcomes of early ablation of Sox2 in the developing telencephalon, via FoxG1-Cre, is the lack of the hippocampal fissure followed by an extreme reduction of the DG. Radial glia scaffold disorganization due to knock-out of the transcription factor Nf1b leads to lack of a specific hippocampal GFAP-positive glial population, lack of hippocampal fissure and DG without affecting cell proliferation, CRC differentiation or Wnt signalling (Barry et al., 2008); this suggests that the loss and disorganization of GFAP-positive cells, seen in our mutants specifically in the developing hippocampus (Fig. 3), might constitute a cellular mechanism contributing to the defective DG development in early Sox2 mutants.

Knock-out of P73 in CH-derived CRC cells leads to lack of hippocampal fissure and DG, as previously mentioned (Meyer et al., 2004). CRC are known to regulate RG formation both in the cortex and in the developing hippocampus (Forster et al., 2002; Frotscher, Haas, & Forster, 2003); conversely, RG has been shown to be important for the correct positioning of CRC cells (Kwon, Ma, & Huang, 2011). Our data suggests that Sox2 does not regulate proliferation in the medial telencephalon at E12.5 (Fig. S2); it is possible that it regulates aspects of differentation of RG and CRC.

### Functional alteration of hippocampal circuitry in Sox2-ablated mice

In Emx1-Cre Sox2-deleted mice, we observed functional alterations in the excitatory transmission along the serial transmission pathway of the hippocampal formation, and particularly an inbalance in the excitatory input onto CA3 and CA1 pyramidal neurons (Fig. 7). Considering that i) the main effects of Sox2 ablation are produced during hippocampal embryogenesis, ii) Sox2 is not expressed in CA3/CA1 neurons, iii) Sox2 deletion caused negligible alterations in pyramidal neuron excitability and excitatory synaptic efficacy, we attribute most of the observed functional effects to altered maturation of the connectivity pattern of hippocampal formation. Neural circuits in the hippocampal formation comprise both serial and parallel pathways. DG is regulated by cortical input from entorhinal layer III, and projects to CA3. However, entorhinal layer III also projects to CA3. Moreover, CA3 displays profuse recurrent reciprocal connections between pyramidal neurons (Witter & Amaral, 2004). Therefore, the higher EPSC frequency we observed in CA3 pyramidal neurons of Sox2-deleted mice could be caused by: i) a denser innervation from entorhinal layer III, permitted by the lower entorhinal input to the hypoplastic DG; ii) an increased recurrent collateral connectivity between CA3 cells, fostered by the absence of the physiological stimulus from DG; iii) a decreased recurrent inhibition on CA3 pyramidal cells, as mossy fibers from DG also regulate GABAergic interneurons in CA3 (Acsady, Kamondi, Sik, Freund, & Buzsaki, 1998). We cannot presently distinguish between these mechanisms, which are not mutually exclusive. Nonetheless, the increased excitatory input we observed in CA3 pyramidal cells is consistent with the epileptic phenotype frequently associated with the brain malformations caused by Sox2 mutations (Sisodiya et al., 2006). Considering the peculiar propensity of CA3 region to develop seizure-like activity (de la Prida, Huberfeld, Cohen, & Miles, 2006; Miles & Wong, 1983), we hypothesize that increased excitatory activity in CA3 of Sox2-deleted mice could facilitate seizure onset, perhaps through CA3 projection to septal areas (Colom, 2006; Swanson & Cowan, 1977).

By contrast, the excitatory input on CA1 pyramidal neurons was lower in Emx1-Cre cKO mice. This could be caused by increased local feed-back inhibition by GABAergic neurons, because of overstimulation by the overactive CA3 fibers. Alternatively, in the absence of proper DG input, the partial disorganization of CA3 connectivity could favor recurrent collaterals at the expense of Schaffer collaterals. Regardless of the specific mechanism, our results demonstrate that Sox2 ablation at early developmental stages unbalances the normal CA3 to CA1 excitatory input, which could contribute to explain some of the cognitive alterations observed in Sox2 mutants. Although early Sox2 ablation leads to severe DG hypoplasia, many cognitive functions can be carried out even when hippocampal volume is strongly reduced (Moser & Moser, 1998; Sisodiya et al., 2006). It is therefore not surprising that the effects of Sox2 ablation on cognition of viable animals are subtle. Nonetheless, evidence is available in humans about a variety of cognitive alterations associated with Sox2 mutations (Ragge et al., 2005; Sisodiya et al., 2006). In general, CA1 is the main output channel of the hippocampal formation, and is thought to compare the entorhinal cortex input (conveying the present state of the environment) with the CA3 input (conveying mnemonic representations of expected events based on external signals; (Knierim & Neunuebel, 2016). Our results suggest that Sox2 malfunction may cause cognitive damage by altering such comparative function of CA1.

### Conclusion and perspective

Overall, our work shows that Sox2 controls (directly, or indirectly) the activity of multiple, functionally interconnected genes, forming a gene regulatory program active and required at very early stages of hippocampal development. Reduced activity of this program leads to essentially absent (FoxG1-Cre mutants) or reduced (Emx1-Cre mutants) development of the hippocampus, in particular the DG. In the Emx1 mutants, which are viable, hippocampal physiology is importantly perturbed. These findings may provide novel perspectives for therapy approaches of genetic brain disease rooted in defective hippocampal development.

## Materials and methods

### Mouse strains

Mutant mice were obtained by crossing the Sox2Flox (Favaro et al., 2009) line with the following lines: FoxG1-Cre (Hébert & McConnell, 2000), Emx1-Cre (Gorski et al., 2002) and Nestin-Cre; Sox2ßGeo (Tronche et al., 1999);Medina, 2004 #506;Favaro, 2009 #3}. The mouse line Sox2-CreERT2 (Favaro et al., 2009), was crossed to a transgenic mouse line with a *loxP-EYFP* reporter of Cre activity (Rosa26R-EYFP) (Srinivas et al., 2001) to determine the progeny of Sox2 expressing cells following tamoxifen injection (as described below).

The day of vaginal plug was defined as embryonic day 0 (E0) and the day of birth as postnatal day 0 (P0).

Genotyping of adult mice or embryos was performed with the following primers:

Sox2 Flox Forward: 5’AAGGTACTGGGAAGGGACATTT 3’
Sox2 Flox Reverse: 5’AGGCTGAGTCGGGTCAATTA 3’
FoxG1-Cre Forward: 5’ AGTATTGTTTTGCCAAGTTCTAAT 3’
FoxG1-Cre Reverse: 5’AGTATTGTTTTGCCAAGTTCTAAT 3’
Emx1-Cre IRES Forward: 5’AGGAATGCAAGGTCTGTTGAAT 3’
Emx1-Cre IRES Reverse: 5’ TTTTTCAAAGGAAAACCACGTC 3’
Nestin-Cre Forward: 5’ CGCTTCCGCTGGGTCACTGTCG 3’
Nestin-Cre Reverse: 5’ TCGTTGCATCGACCGGTAATGCAGGC 3’
R26R-EYFP Forward: 5’ TTCCCGCACTAACCTAATGG 3’
R26R-EYFP Reverse: 5’ GAACTTCAGGGTCAGCTTGC 3’
Sox2-CreERT2 Forward: 5’ TGATCCTACCAGACCCTTCAGT 3’
Sox2-CreERT2 Reverse: 5’ TCTACACATTTTCCCTGGTTCC 3’

The FoxG1-Cre mouse line was maintained in 129 background as recommended in (Hébert & McConnell, 2000). The other mouse lines were maintained in a mixed background enriched in C57BL/6 and DBA.

All procedures were in accordance with the European Communities Council Directive (2010/63/EU and 86/609/EEC), the National Institutes of Health guidelines, and the Italian Law for Care and Use of Experimental Animals (DL26/14). They were approved by the Italian Ministry of Health and the Bioethical Committees of the University of Milan-Bicocca.

### In situ hybridization

*In situ* hybridization was performed essentially as in (Mercurio et al., 2019). Briefly, embryonic brains and P0 brains were dissected and fixed overnight (O/N) in paraformaldehyde 4% in PBS (Phosphate Buffered Saline; PFA 4%) at 4°C. The fixed tissue was cryoprotected in a series of sucrose solutions in PBS (15%, 30%) and then embedded in OCT (Killik, Bio-Optica) and stored at −80°C. Brains were sectioned (20 μm) with a cryostat, placed on a slide (Super Frost Plus 09-OPLUS, Menzel) and stored at −80°C. Slides were then defrosted, fixed in formaldehyde 4% in PBS for 10 minutes (min), washed 3 times for 5 min in PBS, incubated for 10 min in acetylation solution (for 200 ml: 2.66 ml triethanolamine, 0.32 ml HCl 37%, 0.5 ml acetic anhydride 98%) with constant stirring and then washed 3 times for 5 min in PBS. Slides were placed in a humid chamber and covered with prehybridization solution (50% formamide, 5X SSC, 0.25 mg/ml tRNA, 5X Denhardt’s, 0.5 μg/ml salmon sperm) for at least 2 hours (h) and then incubated in hybridization solution (fresh prehybridization solution containing the digoxygenin (DIG)-labelled RNA probe of interest) O/N at 65°C. Slides were washed 5 min in 5X SSC, incubated 2 times in 0.2X SSC for 30 min at 65°C, washed 5 min in 0.2X SSC at room temperature (RT) and then 5 min in Maleic Acid Buffer (MAB, 100 mM maleic acid, 150 mM NaCl pH 7.5). The slides were incubated in blocking solution (10% sheep serum, 2% blocking reagent (Roche), 0.3% Tween-20 in MAB) for at least 1 h at RT, then covered with fresh blocking solution containing anti-DIG antibody Roche © 1:2000 and finally placed O/N at 4°C. Slides were washed in MAB 3 times for 5 min, in NTMT solution (100 mM NaCl, 100 mM Tris-HCl pH 9.5, 50 mM MgCl2, 0.1% Tween-20) 2 times for 10 min and then placed in a humid chamber, covered with BM Purple (Roche), incubated at 37°C until desired staining was obtained (1-6 h), washed in water for 5 min, air dried and mounted with Eukitt (Sigma).

The following DIG-labelled probes were used: *Sox2* (Avilion et al., 2003), *Cadherin8* (Korematsu & Redies, 1997), *Tbr2* (Bulfone et al., 1999), *Reelin* (a gift from Luca Muzio, HSR Milan), *Cxcr4* (Lu et al., 2002), *Cxcl12* (Lu et al., 2002), *Wnt3A* (Grove et al., 1998), *Wnt2b* (Grove et al., 1998), *Wnt5a* (Grove et al., 1998), *Gli3* (a gift from Luca Muzio, HSR Milan), *Lhx2* (a gift from Shubha Tole, Tata Institute Mumbai), *P73* (a gift from Olivia Hanley, UZH). The *P73* probe was transcribed directly from a PCR product, obtained from E12.5 cDNA, with the following primers: Forward 5’ AGCAGCAGCTCCTACAGAGG 5’ and Reverse 5’ <u>TAATACGACTCACTATAGGG</u>CCTTGGGAAGTGAAGCACTC 3’ (which includes the T7 promoter underlined).

### Immunohistochemistry

Immunohistochemistry was performed essentially as in (Cerrato et al., 2018). Brains were dissected, fixed, embedded and sectioned as for *in situ* hybridization, except for fixation in PFA4% that was often 3-4h at 4°C. Sections were washed in PBS 5 min, unmasked in citrate buffer (Na Citrate 0.01M, Citric acid 0.01M pH6) by boiling in a microwave 3 min and then washed in PBS 10 min at RT. Sections were blocked with blocking solution (FBS 10%, Triton 0.3%, PBS1X) for 1h at RT, then incubated O/N in blocking solution with primary antibodies: anti-mSOX2 (R&D Systems MA2018, 1:50), anti-P73 (Neomarkers, 1:150), anti-Reelin (Millipore MAB5364, 1:500), anti-Tuj1 (Covance, 1:400), anti-GFP (Invitrogen A10262, 1:500, used to detect EYFP expressing cells), anti-GFAP (Dako, 1:500). Slides were then washed in PBS 2 times, 10 min each, and incubated in blocking solution containing the secondary fluorescent antibody (1:1000, Alexa Fluor Invitrogen) for 1h 30 minutes at RT. Slides were then washed in PBS twice, 10 minutes each, and then mounted with Fluormount (F4680, Sigma) with 4’,6-diamidino-2-phenylindole (DAPI) and imaged with a confocal microscope (Nikon A1R) and with a Zeiss Axioplan 2 Fluorescent microscope for anti-GFAP immunostainings.

### Lineage tracing of progeny of Sox2 expressing progenitors

R26R-EYFP females were crossed with Sox2-CreERT2 males. E9.5 pregnant females were injected intraperitoneally with tamoxifen (20 mg ml^-1^ in ethanol/corn oil 1:10, 0.1 mg per g of body weight) that induces Cre recombinase activity in the Sox2 telencephalic expression domain (Favaro et al., 2009) and therefore turns on EYFP in this expression domain. Embryos were collected at E15.5, fixed in 4% PFA O/N, embedded in OCT and sectioned at the cryostat (20-μm sections) as for *in situ* hybridization (see above).

### EdU tracing

Ethynyldeoxyuridine (EdU, Molecular Probes) was injected in E12.5 pregnant females at 50 μg/g body weight. Embryos were collected 30 min after injection, fixed O/N in PFA 4% and embedded for cryostat sectioning as above. Edu incorporation was detected on sections (20 μm) with the Click-iT EdU Kit Alexa Fluor 594 (C10354, Thermo Fisher) following manufacturer’s instructions. Briefly, slides were washed twice in PBS 2 min each and incubated for 20 min at RT in Triton 0.5% in PBS. Slides were then washed in Triton 0.1% in PBS 3 times, 3 min each. Sections were incubated 30 min in the dark with EdU Click reaction according to manufacturer’s instructions. Slides were then washed in PBS 3 times 5 min, stained with DAPI, mounted with Fluoromount (F4680, Sigma) and imaged with a confocal microscope (Nikon A1R). The number of EdU positive cells in the cortical hem and dentate neuroephitelium was counted on at least 3 consecutive coronal sections for each brain. Data are represented as mean ±standard deviation.

### Brain slices

For patch-clamp experiments, coronal sections (300 μm thick) containing the hippocampal region (1.22 mm to −2.70 mm from bregma) were prepared from mice of both sexes (6M and 10F) aged P19-P31, by applying standard procedures (Aracri, Meneghini, Coatti, Amadeo, & Becchetti, 2017).

### Patch-clamp recording and data analysis

Cells were examined with an Eclipse E600FN direct microscope, equipped with water immersion DIC objective (Nikon Instruments, Milano, Italy), and digital CCD C8484-05G01 IR camera with HCImage Live acquisition software (Hamamatsu Photonics Italia, Arese, Italy). Stimulation and recording were carried out in whole-cell mode, by using a Multiclamp 700A amplifier (Molecular Devices, Sunnyvale, CA), at 33-34°C. Borosilicate capillaries (OD 1.5 mm; Corning Inc., NY) were pulled (2-3 MΩ) with a Flaming/Brown P-97 micropipette puller (Sutter Instruments, Novato, CA). Series resistance after patch rupture was usually around 10-15 MΩ and was compensated up to at least 70%. Cell capacitance was also compensated. Synaptic currents and action potentials were low-pass filtered a 2 kHz e digitized at 5 kHz with Digidata 1322A / pClamp 9.2 (Molecular Devices). During recording, slices were perfused (∼2 ml/min) with artificial cerebrospinal fluid, containing (mM): 135 NaCl, 21 NaHCO_3_, 0.6 CaCl_2_, 3 KCl, 1.25 NaH_2_PO_4_, 1.8 MgSO_4_, 10 D-glucose, aerated with 95% O_2_ and 5% CO_2_ (pH 7.4). Pipette contained (mM): 140 K-gluconate, 5 KCl, 1 MgCl_2_, 0.5 BAPTA, 1 MgATP, 0.3 NaGTP, 10 HEPES (pH 7.26). Resting membrane potential (V_rest_) was determined in open circuit mode (I=0), immediately after reaching the whole-cell configuration. No correction was applied for liquid junction potentials. Series resistance was monitored throughout the experiment by applying small stimuli around V_rest_. Cells were discarded when Rs was higher than 15 MΩ.

Action potentials and EPSCs were analysed with Clampfit 9.2 (Molecular Devices), MiniAnalysis, and OriginPro 9.1(OriginLab Corporation, Northampton, MA, USA), as previously reported (Aracri, Amadeo, Pasini, Fascio, & Becchetti, 2013; Aracri et al., 2017).

### AZD and LiCl treatment

AZD 1080 (Axon Medchem, Axon Catalog ID: 2171) diluted in ascorbic acid 0.5%/EDTA 0.01%, was administered to pregnant females once a day by oral gavage from E9.5 to E12.5. We administered 5 μl AZD/g of body weight (AZD 0.375 μg/μl at E9.5 and E10.5, AZD 0.75 μg/μl at E11.5 and E12.5). Embryos were then collected at E14.5. Ascorbic acid 0.5%/EDTA 0.01% was administered as a control.

LiCl, or NaCl as a control, were injected intraperitoneally in pregnant female from E9.5 to E14.5 or from E10.5 to E12.5 once a day at the same time. No difference was observed between the two injection time windows. 10 μl/ g of body weight of 600 mM LiCl or 600 mM NaCl were injected. Embryos were collected at E18.5 and processed for *in situ* hybridization. Injection of AZD 1080 or LiCl in pregnant females at E8.5 led to abortions.

The number of *Reelin*-positive cells at the hippocampal fissure and in the cortex was counted using Photoshop CC 2015 on five consecutive coronal sections of each brain. Data are represented as mean ± standard deviation and were statistically analyzed using unpaired Student’s T-test, ***p<0.005.

### Luciferase constructs

The DNA region in the Gli3 second intron overlapping the SOX2 peak, and corresponding to the VISTA enhancer (coordinates under the embryo in Fig. 6A) was PCR-amplified from the vector where it had been cloned upstream to the lacZ reporter (a gift from T. Theil; (Hasenpusch-Theil et al., 2012)), and cloned upstream to the tk promoter in the Tk-luc vector (Mariani et al., 2012), into the KpnI and NheI restriction sites.

### Transfection experiments

The transfection experiments were performed essentially as previously described (Mercurio et al., 2019; Panaliappan et al., 2018). In particular, Neuro-2a cells were plated in Minimal Essential Medium Eagle (MEM; SIGMA), supplemented with 10% Foetal Bovine Serum, L-Glutamine, penicillin and streptomycin. For transfection, cells were plated in 12-well plates at 1.5×10^5^ cells/well, and transfected on the following day using Lipofectamine 2000 (Invitrogen). Briefly, medium in each well was replaced with 1 ml of MEM medium (with no addition) mixed with 2 μl of Lipofectamine 2000, and DNA. After 4 hours from transfection, the medium was replaced with complete medium. A fixed amount of 300 ng of luciferase reporter plasmid was used for each well, with increasing amounts of Sox2 expressing vector (Favaro et al., 2009; Mariani et al., 2012), or the corresponding control “empty” vector (not containing the transcription factor’s cDNA), in the following luciferase vector:expressing vector molar ratios (indicated in Fig.6): +, 1:0.050; ++, 1:0.075; +++, 1:0.125; ++++, 1:0.25; +++++, 1:0.5. The pBluescript vector was added to transfection DNA to equalize the total amount of transfected DNA to a total of 800 ng for each reaction. After 24 hours, total cellular extracts were prepared, and Luciferase activity was measured with a Promega Luciferase Assay System, according to the manufacturer’s instructions.

---

**Supplementary Figure 1.**
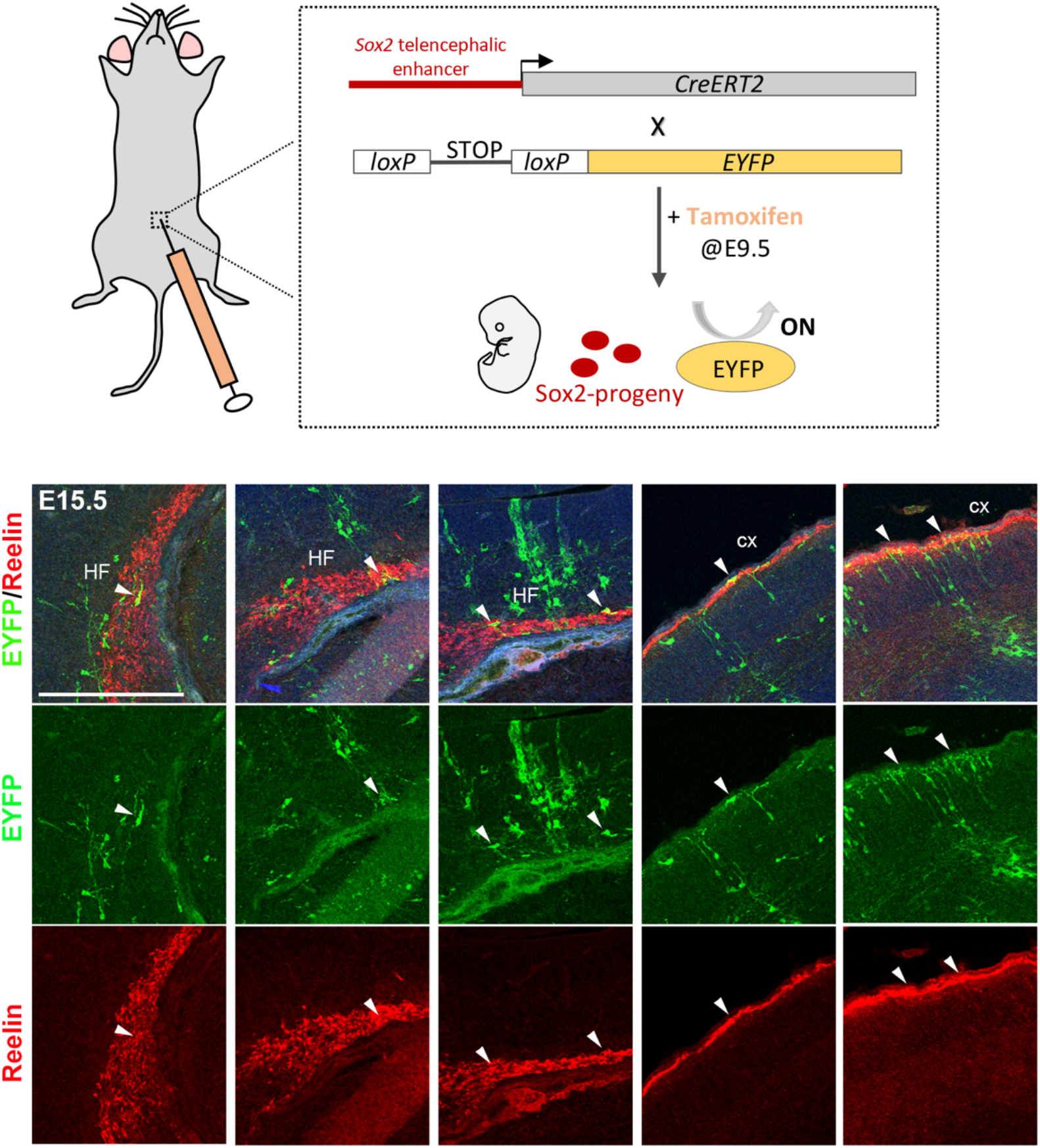
Lineage tracing of Sox2 expressing progeny in CR cells in the hippocampal fissure (HF) and the cortex. Top, scheme depicting the mouse crosses used to trace the progeny of Sox2 expressing cells. A mouse in which expression of an inducible Cre recombinase (cre-ERT2) was under the control of a Sox2 telencephalic enhancer was crossed to a mouse in which YFP expression would be turned on when Cre recombinase was expressed, following tamoxifen injection. Bottom, immunofluorescence for GFP (green) and Reelin (red) on coronal sections of brains at E15.5 from pregnant females injected with tamoxifen at E9.5. Arrows indicate cells that are positive for both GFP and Reelin in the hippocampal fissure (HF) and the cortex (cx). Scale bar 200 μm.

**Supplementary Figure 2.**
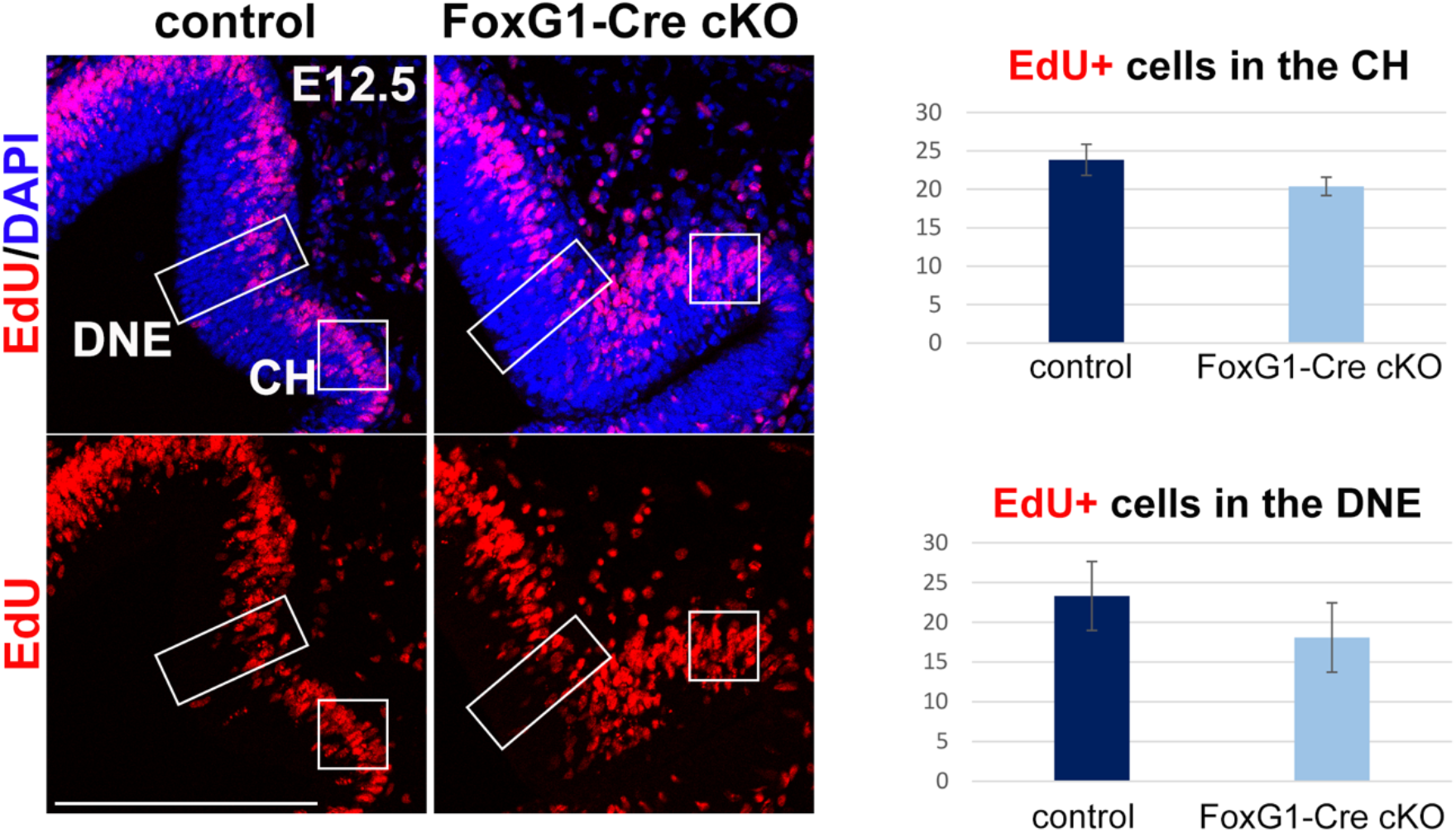
Sox2 early ablation does not appear to affect cellular proliferation neither in the cortical hem nor in the dentate neuroepithelium. EdU staining on coronal sections of control and FoxG1-Cre cKO mice at E12.5 injected with EdU and sacrificed 30 minutes later. The graphs on the right show no significant difference in the number of proliferating cells in the cortical hem (CH) and in the dentate neuroephitelium (DNE) at the developmental stage analysed. Data are represented as mean ±standard deviation (controls n=3, mutants n=3). Scale bar 200 μm.

**Supplementary Figure 3.**
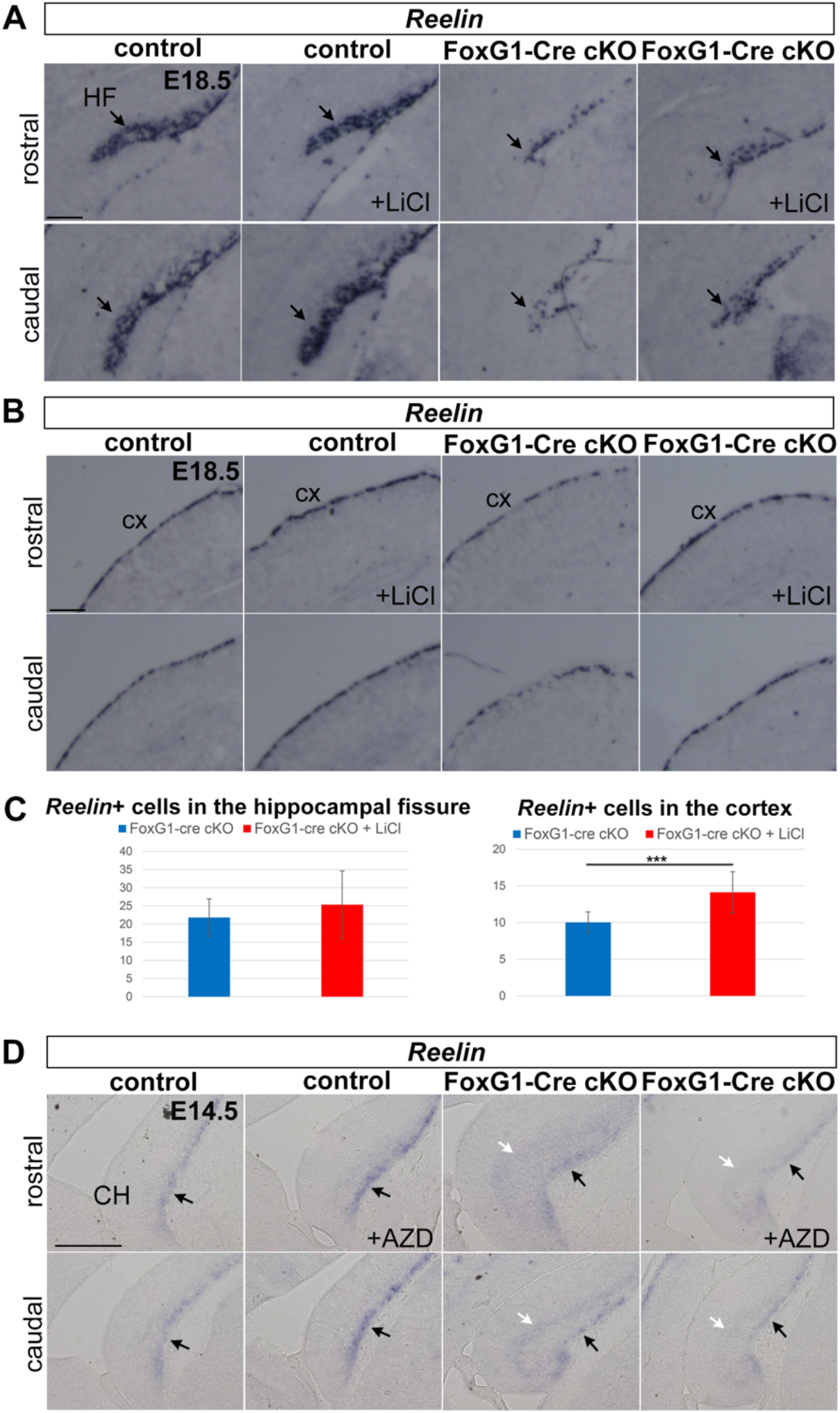
Administration of agonists of the Wnt pathway LiCl and AZD 1080 partially rescues the deficit of C-R cells in the cortex, but only slightly in the hippocampal fissure. *Reelin in situ* hybridization on coronal sections of control or FoxG1-Cre cKO brains injected with Wnt agonists, at the times indicated, and analyzed at E18.5 in the case of LiCl injection **(A,B)** and at E14.5 for AZD 1080 injection **(D)**. LiCl or NaCl were intraperitoneally injected once a day in pregnant females from E10.5 to E12.5. AZD 1080 was administered by oral gavage once a day in pregnant females from E9.5 to E12.5. At least 3 controls and 3 mutants, both treated and untreated, were analyzed. The graphs in **(C)** indicate a significant rescue in the number of C-R cells in the cortex following LiCl injection (*** p<0.005, unpaired Student’s T-test. Mutants untreated n=5, mutants LiCl treated n=5). Error bars represent standard deviation. (D) Black arrows indicate Reelin expressing cells towards the pial side of the cortical hem, white arrows indicate Reelin expressing cells retained in the cortical hem that did not move towards the pia. AZD 1080 treatment rescued this retention. Scale bars 200 μm.

**Supplementary Table 1.**
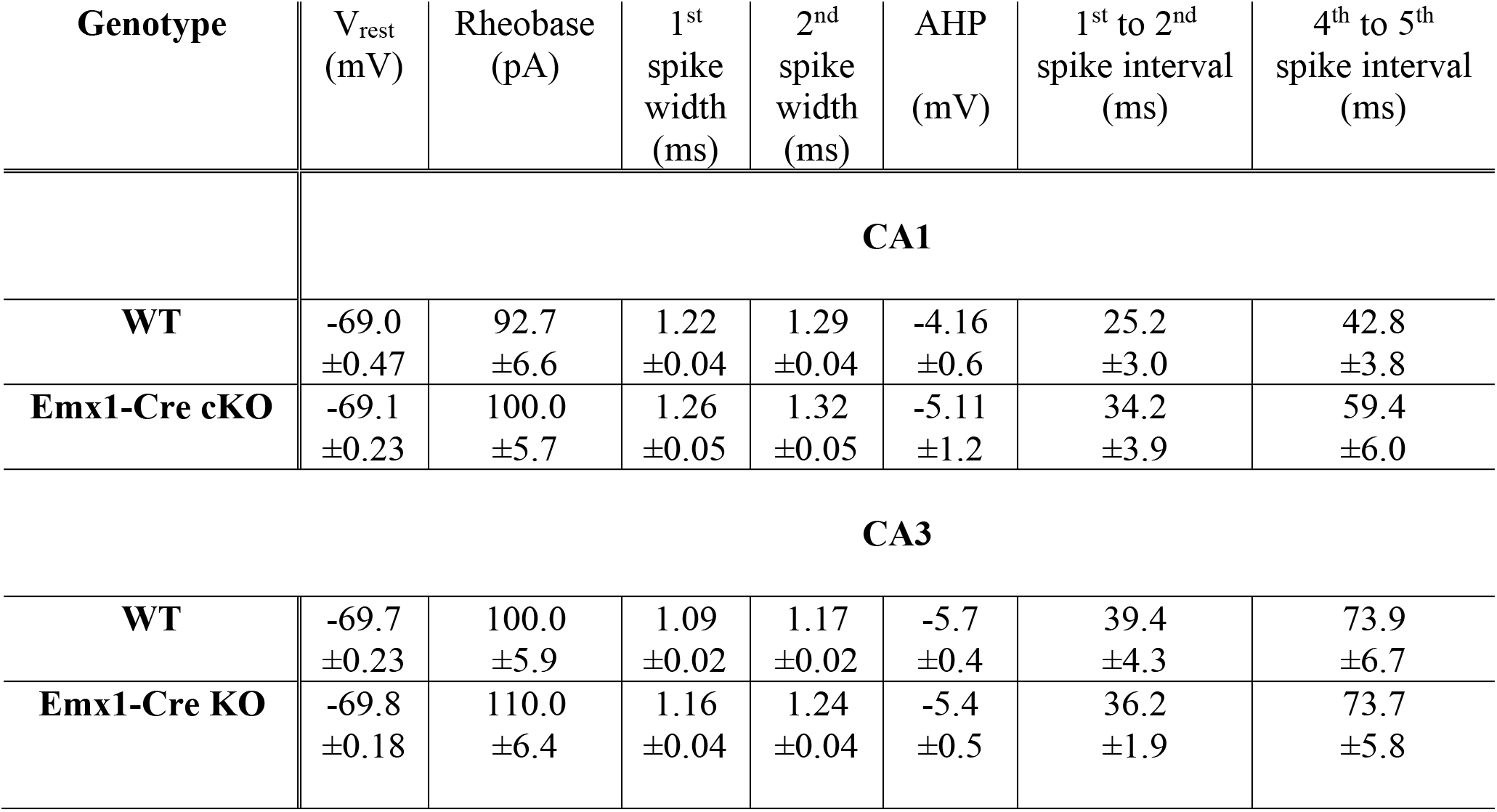
Excitability features of CA1/CA3 pyramidal neurons of Emx1-Cre cKO and control mice. For the WT and Emx1-Cre cKO mice, the main excitability features are given for CA3 and CA1 pyramidal neurons, as indicated. The table reports average V_rest_, rheobase, spike widths (measured at half-amplitude) of the first and second action potential, and AHP amplitude (with respect to V_rest_). As a measure of firing adaptation, the time intervals are reported between the first and second spike and the fourth and fifth spike.

| Genotype | V<sub>rest</sub><br>(mV) | Rheobase<br>(pA) | 1<sup>st</sup><br>spike<br>width<br>(ms) | 2<sup>nd</sup><br>spike<br>width<br>(ms) | AHP<br>(mV) | 1<sup>st</sup> to 2<sup>nd</sup><br>spike interval<br>(ms) | 4<sup>th</sup> to 5<sup>th</sup><br>spike interval<br>(ms) |
| --- | --- | --- | --- | --- | --- | --- | --- |
| **CA1** |  |  |  |  |  |  |  |
| **WT** | -69.0<br>±0.47 | 92.7<br>±6.6 | 1.22<br>±0.04 | 1.29<br>±0.04 | -4.16<br>±0.6 | 25.2<br>±3.0 | 42.8<br>±3.8 |
| **Emx1-Cre cKO** | -69.1<br>±0.23 | 100.0<br>±5.7 | 1.26<br>±0.05 | 1.32<br>±0.05 | -5.11<br>±1.2 | 34.2<br>±3.9 | 59.4<br>±6.0 |
| **CA3** |  |  |  |  |  |  |  |
| **WT** | -69.7<br>±0.23 | 100.0<br>±5.9 | 1.09<br>±0.02 | 1.17<br>±0.02 | -5.7<br>±0.4 | 39.4<br>±4.3 | 73.9<br>±6.7 |
| **Emx1-Cre KO** | -69.8<br>±0.18 | 110.0<br>±6.4 | 1.16<br>±0.04 | 1.24<br>±0.04 | -5.4<br>±0.5 | 36.2<br>±1.9 | 73.7<br>±5.8 |

